# Impaired succinate oxidation prevents growth and influences drug susceptibility in *Mycobacterium tuberculosis*

**DOI:** 10.1101/2021.06.21.449354

**Authors:** Cara Adolph, Matthew B. McNeil, Gregory M. Cook

## Abstract

Succinate is a major focal point in mycobacterial metabolism and respiration, serving as both an intermediate of the TCA cycle and a direct electron donor for the respiratory chain. *Mycobacterium tuberculosis* encodes multiple enzymes predicted to be capable of catalyzing the oxidation of succinate to fumarate, including two different succinate dehydrogenases (Sdh1 and Sdh2) and a separate fumarate reductase (Frd) with possible bi-directional behavior. Previous attempts to investigate the essentiality of succinate oxidation in *M. tuberculosis* have relied on the use of single-gene deletion mutants, raising the possibility that the remaining enzymes could catalyze succinate oxidation in the absence of the other. To address this, we report on the use of mycobacterial CRISPR interference (CRISPRi) to construct single, double, and triple transcriptional knockdowns of *sdhA1*, *sdhA2*, and *frdA* in *M. tuberculosis*. We show that the simultaneous knockdown of *sdhA1* + *sdhA2* is required to prevent succinate oxidation and overcome the functional redundancy within these enzymes. Succinate oxidation was demonstrated to be essential for the optimal growth of *M. tuberculosis*, with the combined knockdown of *sdhA1* + *sdhA2* significantly impairing the activity of the respiratory chain and preventing growth on a range of carbon sources. Moreover, impaired succinate oxidation was shown to influence the activity of several antitubercular drugs against *M. tuberculosis*, including potentiating the activity of bioenergetic inhibitors and attenuating the activity of cell wall inhibitors. Together, these data provide fundamental insights into mycobacterial physiology, energy metabolism, and antimicrobial susceptibility.

**Importance:** New drugs are urgently required to combat the tuberculosis epidemic that claims 1.5 million lives annually. Inhibitors of mycobacterial energy metabolism have shown significant promise clinically; however, further advancing this nascent target space requires a more fundamental understanding of the respiratory enzymes and pathways used by *Mycobacterium tuberculosis*. Succinate is a major focal point in mycobacterial metabolism and respiration; yet the essentiality of succinate oxidation, and the consequences of inhibiting this process, are poorly defined. In this study, we demonstrate that impaired succinate oxidation prevents the optimal growth of *M. tuberculosis* on a range of carbon sources and significantly reduces the activity of the electron transport chain. Moreover, we show that impaired succinate oxidation both positively and negatively influences the activity of a variety of anti-tuberculosis drugs. Combined, these findings provide fundamental insights into mycobacterial physiology and drug susceptibility that will be useful in the continued development of bioenergetic inhibitors.

## Introduction

Tuberculosis (TB) is a leading cause of infectious disease morbidity and mortality globally, with 10 million new cases and 1.4 million deaths reported in 2019 (1). Successful treatment of infections with *Mycobacterium tuberculosis*, the causative agent of the disease, requires six months of combination therapy involving two months of intensive treatment with ethambutol (EMB), isoniazid (INH), rifampicin (RIF) and pyrazinamide (PZA), followed by a further two months of treatment with INH and RIF (2). Infections with multidrug-resistant (MDR) and extensively drug-resistant (XDR) strains of *M. tuberculosis* require up to two years of chemotherapy with a combination of 8-10 antibiotics and have cure rates as low as 39% for XDR-TB infections (1). Thus, global TB control urgently depends on the development of new drugs and regimens that can quickly and effectively treat both drug-sensitive and drug-resistant TB infections.

Mycobacterial energy generation has emerged as a promising target space for antitubercular drug development. Enzymes within the central carbon metabolism of *M. tuberculosis* are increasingly being recognized as essential mediators of pathogenicity and their inhibition or deletion is often bactericidal *in vivo* (3–9). Likewise, a functional respiratory chain is essential for the viability of both replicating and non-replicating *M. tuberculosis* (10–14). Several inhibitors of the mycobacterial respiratory chain have now been identified including the only three new TB drugs to be approved in the last 50 years; bedaquiline (BDQ), which inhibits the mycobacterial F1Fo ATP synthase (15–17) and pretomanid (PA-824) and delamanid, which result in NO-induced respiratory poisoning (18–22). The clinical efficacy of these respiratory inhibitors (23–26), especially when used in combination (i.e., the Nix-TB regimen (27)), demonstrates the significant treatment shortening potential of targeting mycobacterial energy generation.

Succinate oxidation is a major focal point in mycobacterial energy generation and directly couples central carbon metabolism to the respiratory chain (28). The oxidation of succinate to fumarate is catalyzed by succinate dehydrogenase (SDH) enzymes, which couple succinate oxidation in the TCA cycle to the reduction of menaquinone in the electron transport chain (ETC) (succinate + menaquinone ↔ fumarate + menaquinol) (29, 30). Fumarate reductase (FRD) enzymes catalyze the reverse reaction: fumarate reduction coupled to menaquinol oxidation. SDH and FRD enzymes encoded by bacteria are highly similar and are often functionally interchangeable (30, 31). *M. tuberculosis* encodes three different SDH / FRD enzymes which are all precited to be capable of catalyzing the oxidation of succinate to fumarate (32). This includes two different succinate dehydrogenase enzymes (Sdh1; Rv0249c-Rv0247c and Sdh2; Rv3316- Rv3319), as well as a separate fumarate reductase with possible bi-directional behavior (Frd; Rv1552-Rv1555) (32). All three enzymes have distinct phylogenies, prosthetic groups and predicted biochemistries (30, 33–36), and are differentially expressed in *M. tuberculosis* (37–39), suggesting that these enzymes perform distinct, but overlapping, roles.

Previous attempts to elucidate the function of these individual enzymes in *M. tuberculosis* have focused on the characterization of single-gene deletion mutants (37, 40). However, this leaves the possibility that the remaining enzymes can carry out succinate oxidation in the absence of the other. Consistent with this, single deletion mutants in *M. tuberculosis* only have minor (i.e., for Δs*dh1* and Δ*sdh2*) (40) or no (i.e., for Δ*frd*) (37) phenotypes, while the construction of double deletion mutants was unsuccessful (40). Likewise, while high-throughput transposon hybridization (TraSH) screens have hinted at the essentiality of SDH enzymes for mycobacterial growth and persistence (3, 41, 42), they have also been unable to overcome the functional redundancy in SDH catalysis. Consequently, the essentiality of succinate oxidation in *M. tuberculosis* is incompletely understood.

Here, we used mycobacterial CRISPR Interference (CRISPRi) (43–45) to transcriptionally repress the expression of *sdhA1*, *sdhA2*, and *frdA* in *M. tuberculosis* alone and in combination, allowing us to overcome the functional redundancy within succinate oxidation. We demonstrate that Sdh1 is the primary aerobic SDH in *M. tuberculosis*, although its loss can be compensated for by Sdh2 but not Frd, which appears to be unable to catalyze succinate oxidation. Succinate oxidation was essential for the optimal growth of *M. tuberculosis*, with the simultaneous knockdown of *sdhA1* + *sdhA2* significantly impairing the activity of the respiratory chain and preventing growth on a range of carbon sources. Moreover, we demonstrate that impaired succinate oxidation both positively and negatively affects the susceptibility of *M. tuberculosis* to a variety of anti-TB drugs.

## Results

### Succinate oxidation is essential for the optimal growth of *M. tuberculosis*

To elucidate the roles and essentialities of the SDH and FRD enzymes in *M. tuberculosis*, we used CRISPR Interference (43) to knockdown the expression of Sdh1, Sdh2 and Frd alone and in combination. Three guide RNAs (i.e., sgRNA) were designed to target the catalytic A subunit of Sdh1, Sdh2 and Frd (Figure 1A-C), which resulted in high-level transcriptional repression (50 to 100-fold) of target genes in single and multiplexed constructs (Figure 1D and E). Interestingly, expression of the sgRNA targeting *sdhA1* also resulted in reduced expression of *frdA* in multiplexed repression constructs (Figure 1.E), potentially suggesting a shared regulatory mechanism governing Sdh1 and Frd gene expression. Transcriptional repression of either *sdhA1*, *sdhA2* or *frdA* alone did not affect the growth of *M. tuberculosis* strain mc^2^6230 in 7H9 media supplemented with OADC (Figure 1F), consistent with the previously published single deletion mutants (37, 40). Simultaneous knockdown of both *sdhA1* + *sdhA2*, with or without *frdA*, was required to prevent growth in 7H9-OADC media (Figure 1G) and had bacteriostatic consequences on cell viability (Figure 1H). The combined knockdown of *frdA* with either *sdhA1* or *sdhA2* displayed no growth defect in the same media (Figure 1G).

**Figure 1:**
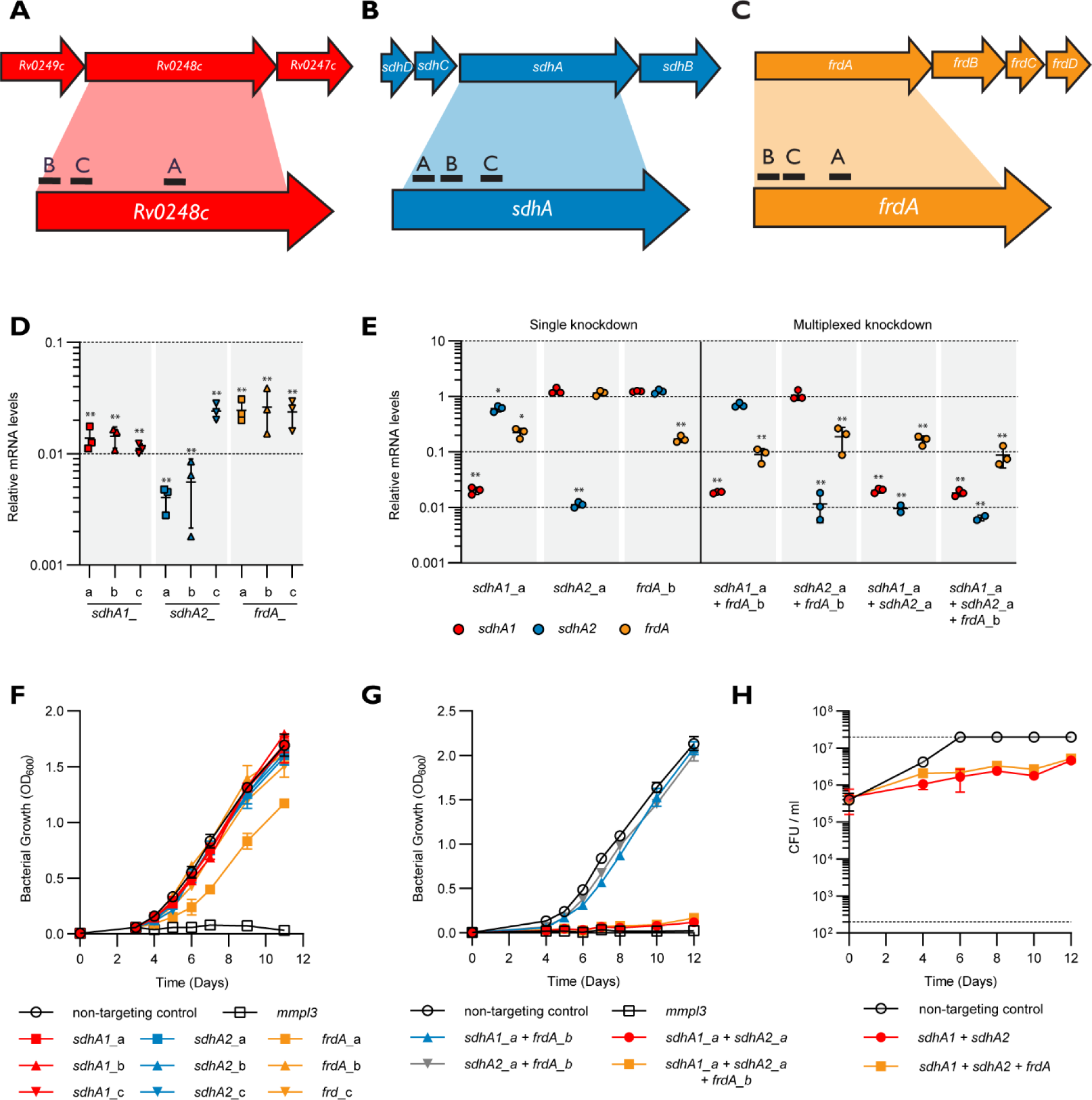
Single and multiplexed transcriptional repression of *sdhA1*, *sdhA2* and *frdA* in *M. tuberculosis* using CRISPRi. **(A-C)** Location of sgRNAs targeting the catalytic subunits of Sdh1 **(A)**, Sdh2 **(B)** and Frd **(C)** in *M. tuberculosis*. sgRNAs were co- expressed with a dCas9sth1 under the control of an ATc inducible promoter. **(D)** CRISPRi achieves high-level knockdown of target genes in single repression constructs. RNA was harvested 72 hours after inducing knockdown (100 ng/ml ATc) and quantified by qPCR. mRNA is expressed relative to a strain expressing a non-targeting control. Results are mean ± standard deviation (SD) of three technical triplicates. ** indicates a *p*-value of < 0.005 from a one-way ANOVA with a Dunnet correction comparing each sgRNA to the non-targeting control. **(E)** CRISPRi achieves high-level knockdown of *sdhA1*, *sdhA2* and *frdA* in single, double, and triple gene repression constructs. Gene knockdown was quantified and visualized as in **D**. Statistical significance was calculated using a one-way ANOVA and a Dunnet test for multiple comparisons of the gene expression in each strain against the non-targeting control; * *p* < 0.01, ** *p* < 0.001. **(F)** Consequences of the single knockdown of *sdhA1*, *sdhA2* and *frdA* on the growth of *M. tuberculosis* in 7H9 media supplemented with OADC. Knockdown was induced at time 0hr with ATc (100 ng/ml). A *mmpL3* targeting sgRNA, which has bactericidal consequences on the viability of *M. tuberculosis* (45), was included as a positive inhibition control. The means and standard deviation of three replicates are shown. **(G and H)** Consequences of multiplexed *sdhA1*, *sdhA2* and *frdA* gene repression on the growth **(G)** and viability **(H)** of *M. tuberculosis* in 7H9 media supplemented with OADC. Dashed horizontal lines in **H** represent the upper and lower limits of detection of CFU. The means and standard deviation of three replicates are shown. Data in **G** and **H** are representative of three independent experiments.

To further understand the contribution of each enzyme to mycobacterial growth, we investigated the ability of the *M. tuberculosis* SDH / FRD knockdown strains to grow on a range of fermentable and non-fermentable carbon sources. When provided with succinate as the sole carbon and energy source, the knockdown of *sdhA1* alone, but not *sdhA2* or *frdA*, had a minor (approx. 10%) defect in growth rate (Figure 2A), consistent with its proposed role as the primary aerobic SDH (40). For all single transcriptional knockdowns, no significant difference in growth compared to a non-targeting control was observed in media containing glucose, glycerol, acetate, or a mixture of glucose and acetate (Figure 2B-E). This demonstrates that these enzymes are individually dispensable for the growth of *M. tuberculosis*, regardless of carbon source.

**Figure 2:**
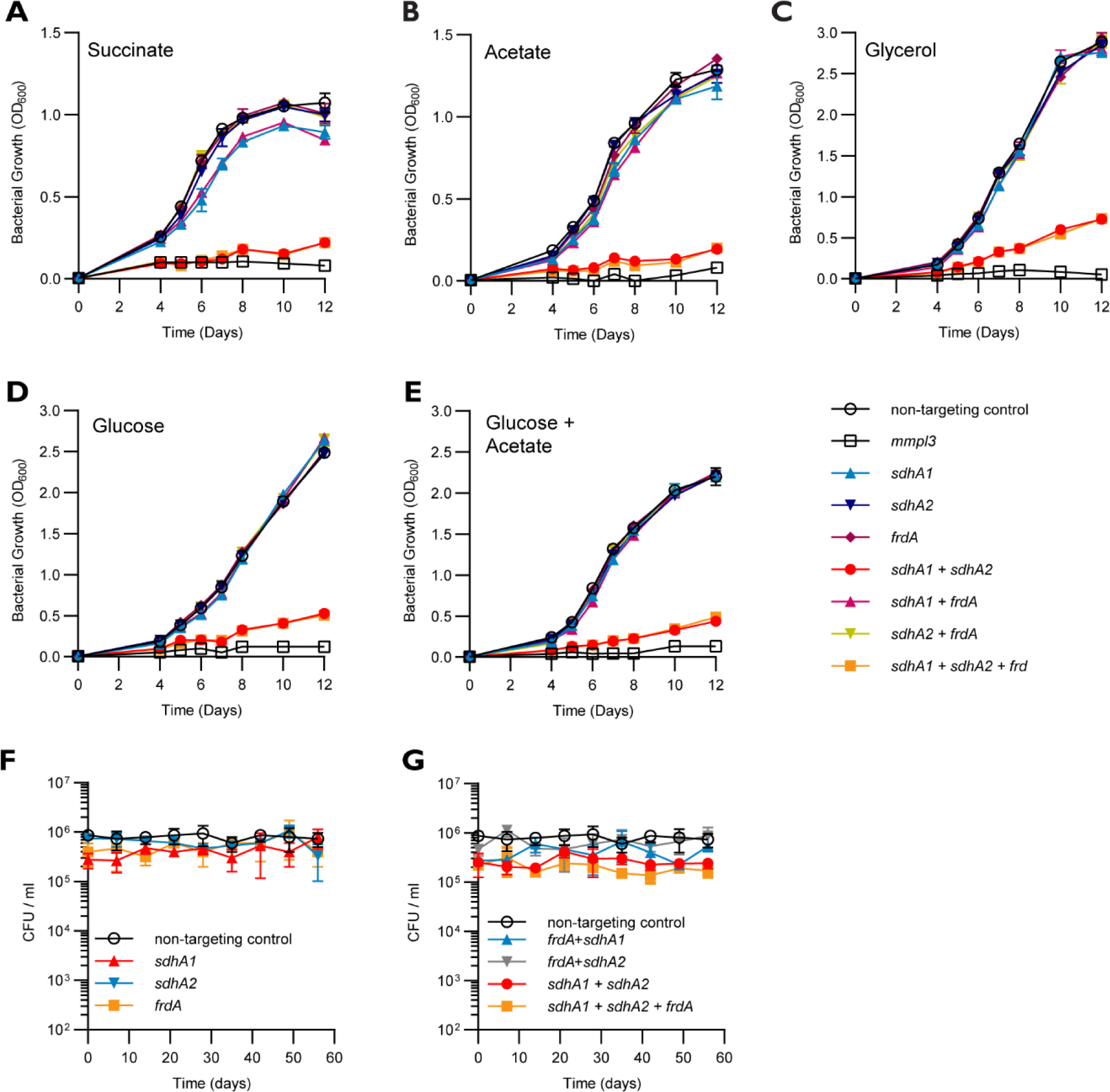
Succinate oxidation is essential for the optimal growth of *M. tuberculosis* but is dispensable for survival under nutrient starvation. **(A – E)** Growth profiles of *M. tuberculosis* single and multiplexed *sdhA1*, *sdhA2* and *frdA* knockdown strains in 7H9 media supplemented with 30 mM succinate **(A)**, 0.2% acetate **(B)**, 0.2% glycerol **(C)**, 20 mM glucose **(D)**, or a combination of 0.1% acetate and 10 mM glucose **(E)**. Cultures were grown in 10 ml volumes from a starting OD600 of 0.005 and gene knockdown was induced at time 0hr with ATc (100 ng/ml). A *mmpl3* knockdown strain was included as a positive inhibition control. The means and standard deviation of three replicates are shown. Each experiment was repeated twice. **(F and G)** Consequences of the single **(F)** or multiplexed **(G)** depletion of Sdh1, Sdh2 or Frd on the survival of *M. tuberculosis* under a nutrient starvation model of non-replicating persistence. Before entering nutrient starvation, cells were pre-depleted of SDH and FRD enzymes by inducing transcriptional repression of target genes for 8 days. Cultures were then harvested, washed twice in PBS and nutrient starved by inoculating into inkwells containing PBS + tyloxapol. Viability was measured by enumerating CFU/ml over 8 weeks. Error bars represent the standard deviation from three replicates.

The combined transcriptional repression of *sdhA1* + *sdhA2* prevented the optimal growth of *M. tuberculosis* across all carbon sources tested (Figure 2A-E). *M. tuberculosis* repressing both *sdhA1* + *sdhA2* was unable to grow with the non-fermentable carbon sources succinate or acetate as the sole carbon and energy source (Figure 2A and B). The growth impairment of the *sdhA1* + *sdhA2* double knockdown strain was partially rescued when using fermentable carbon sources such as glycerol or glucose, although growth was still significantly impaired (Figure 2C and D). Furthermore, glucose was able to partially rescue the growth impairment of the *sdhA1* + *sdhA2* knockdown when *M. tuberculosis* was grown on a combination of glucose and acetate (Figure 2.3E). Frd was unable to compensate for the depletion of *sdhA1* + *sdhA2* with the additional knockdown of *frdA* showing the same level of growth inhibition as the dual depletion of *sdhA1* + *sdhA2* (Figure 2A-E). This suggests that Frd does not contribute to aerobic growth under the conditions tested. Overall, these data demonstrate that a functional SDH enzyme (i.e., Sdh1 or Sdh2) is essential for the optimal growth of *M. tuberculosis*.

### SDH and FRD enzymes are dispensable for the survival of *M. tuberculosis* under nutrient starvation

We next sought to investigate the contribution of SDH and FRD enzymes to the survival of *M. tuberculosis* under nutrient starved non-replicating persistence (38). We first depleted cells of SDH / FRD enzymes by growing strains for eight days in the presence of ATc (100 ng/ml), before placing knockdown strains into nutrient starved conditions (i.e., PBS). Individual depletion of either Sdh1, Sdh2 or Frd did not affect the survival of *M. tuberculosis* over 8 weeks of nutrient starvation (Figure 2F). Moreover, all double and triple transcriptional knockdown strains had no survival defect under nutrient-starved conditions (Figure 2G). Together, this suggests that SDH and FRD enzymes are dispensable for the survival of *M. tuberculosis* under nutrient starvation.

### Impaired succinate oxidation significantly disrupts the activity of the mycobacterial respiratory chain

The oxidation of succinate to fumarate results in two electrons being donated to the ETC (30). Therefore, we sought to determine the bioenergetic consequences of impaired succinate oxidation on the activity of the *M. tuberculosis* respiratory chain. To achieve this, we pre-depleted cells of SDH / FRD enzymes by inducing transcriptional repression for eight days prior to measuring oxygen consumption rates (OCR) as a proxy of respiratory activity. We measured the OCR when cultures were grown with OADC, or with glycerol or succinate as the sole carbon and energy source as these were the growth conditions where the SDH / FRD knockdown strains were the least and most impaired, respectively (Figure 2A and C).

When grown in media supplemented with OADC, only the combined knockdown of *sdhA1* + *sdhA2* was able to overcome the functional redundancy in succinate oxidation and affect the OCR of *M. tuberculosis* (Figure 3A). All other combinations of transcriptional repression did not influence the OCR of *M. tuberculosis* when grown with OADC (Figure 3A). In contrast, with succinate as the sole carbon source, the single knockdown of *sdhA1* alone, or with *frdA*, reduced the OCR of *M. tuberculosis* by approximately 20% (Figure 3B). This is consistent with the growth impairment of these strains when grown on succinate (Figure 2A) and suggests that Sdh1 is the dominant SDH enzyme in *M. tuberculosis*. No significant difference in OCR was observed for the *sdhA1*, or *sdhA1* + *frdA* knockdown strains in media supplemented with glycerol (Figure 3C).

**Figure 3:**
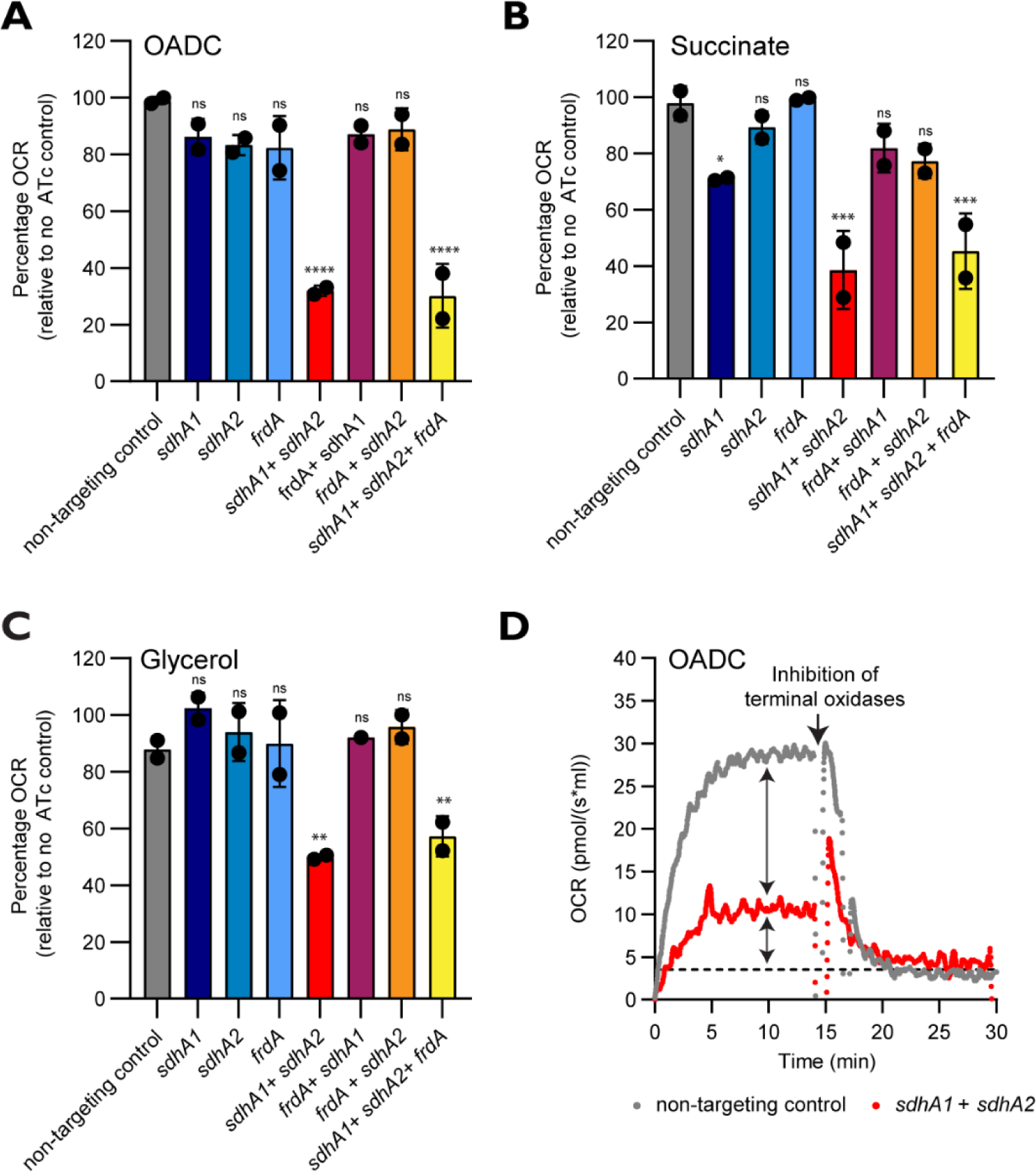
Succinate oxidation is the major contributor of electrons to the respiratory chain in *M. tuberculosis*. **(A - C)** Oxygen consumption rates (OCR) of the single and multiplexed *sdhA1*, *sdhA2* and *frdA* knockdown strains when energized with OADC **(A)**, succinate **(B)** or glycerol **(C)**. Cultures were grown for 8 days in the presence of 100 ng/ml ATc to deplete cells of SDH / FRD enzymes in 7H9 media containing the specified carbon sources before performing OCR measurements on cell suspensions. Data was normalized to the respective no ATc control for each strain. 100% for OADC = ∼30.0 pmol/(s*ml), for succinate = ∼30.0 pmol/(s*ml), and for glycerol = ∼35.0 pmol/(s*ml). Error bars represent the mean and standard deviation of two technical replicates and data is representative of two independent experiments. Statistical significance was calculated using a one-way ANOVA and a Dunnet test for multiple comparisons of each strain against the non-targeting control; ns *p* > 0.05, * *p* < 0.05, ** *p* < 0.01, *** *p* < 0.001 and **** *p* < 0.0001. **(D)** Chemical inhibition of both terminal oxidases with Q203 (400 nM) and ND-011992 (100µM) demonstrates that succinate oxidation is the major contributor of electrons to the respiratory chain. Cultures were grown for 8 days in the presence of 100 ng/ml ATc to deplete cells of SDH / FRD enzymes in 7H9-OADC media before performing OCR measurements on cell suspensions. Dashed horizonal lines in **D** represent the baseline OCR. Arrows represent the contribution of succinate oxidation to the OCR of *M. tuberculosis,* and the residual OCR in the absence of succinate oxidation, respectively. Data are representative of two independent experiments.

The combined repression of both *sdhA1* + *sdhA2* ± *frdA* significantly reduced the respiration rate of *M. tuberculosis* across all three carbon sources tested (Figure 3A-C). The double and triple *sdhA1* + *sdhA2* ± *frdA* knockdown strains had a 50% reduction in OCR compared to their respective no ATc controls in media containing glycerol, and a 70% reduction in OCR when energized with OADC or succinate (Figure 3A-C). On all three carbon sources tested, there was no difference in the OCR between the double *sdhA1* + *sdhA2* knockdown strain and the triple *sdhA1* + *sdhA2* + *frdA* knockdown strain (Figure 3A-C), suggesting that Frd is unable to carry out succinate oxidation under the conditions tested.

To further investigate the contribution of succinate oxidation to total respiratory flux, we used Q203 and ND-011992 to inhibit the activity of both terminal oxidases and abolish respiration (46, 47). The complete inhibition of aerobic respiration highlighted an additional ∼30% respiratory capacity in the endogenous respiration rate of *M. tuberculosis* in the absence of succinate oxidation (Figure 3D). Combined, these results demonstrate that succinate oxidation is the major contributor of electrons to the *M. tuberculosis* respiratory chain, regardless of carbon source (Figure 3A-D).

### Impaired succinate oxidation influences the drug susceptibility of *M. tuberculosis*

TB is treated with multidrug therapies and, as such, any new drug needs to be effective in combination. While bioenergetic inhibitors are proving promising at reducing treatment times, the consequences of inhibiting respiration on the activity of other antibiotics are not clearly understood. Several studies have linked antibiotic killing to the dysregulation of respiration and the production of reactive oxygen species (48–54), while others have demonstrated that inhibiting respiration can attenuate the activity of bactericidal drugs (53, 55–59). Given that the loss of SDH activity reduced the activity of the respiratory chain by approximately 70% (Figure 3), we sought to investigate how the simultaneous knockdown of *sdhA1* + *sdhA2* affected the activity of anti-TB drugs against *M. tuberculosis*. We profiled the *sdhA1* + *sdhA2* double knockdown strain of *M. tuberculosis* against TB drugs with a range of cellular targets including cell wall biosynthesis (INH, PA- 824, ethionamide (ETH), EMB and SQ109), DNA replication (levofloxacin (LEV)), transcription (RIF), protein synthesis (linezolid (LZD) and streptomycin (STREP)), and bioenergetics (BDQ, clofazimine (CFZ), thioridazine (THZ), SQ109, Q203, TB47 and PA- 824). We induced gene knockdown and treated cultures with antibiotics simultaneously on day 0 as previously described (44, 45) and monitored killing (rather than MIC) as a measure of susceptibility as *M. tuberculosis* is significantly impaired for growth when transcriptionally repressing *sdhA1* + *sdhA2* (Figure 1 and 2).

The combined knockdown of *sdhA1* + *sdhA2* resulted in increased killing of *M. tuberculosis* by INH and PA-824 after 10 days of incubation (Figure 4A and B), while the susceptibility to all other drugs tested remained unchanged (Figure S1). We investigated this interaction further by monitoring killing by INH and PA-824 over time. Interestingly, there was no change in the rate of killing by either drug but instead the simultaneous knockdown of *sdhA1* + *sdhA2* prevented the emergence of resistance to both INH and PA-824 (Figure 4C and D). After 3 weeks, no INH or PA-824 resistant mutants were isolated when both Sdh1 and Sdh2 were depleted, whilst INH and PA-824 resistance emerged by day 10 in the no knockdown controls (Figure 4C and D). This could indicate that normal respiratory chain function is required for the development of INH and PA-824 resistance as previously described (14, 60, 61) or it could be the result of the reduced replication rate of *M. tuberculosis* when repressing *sdhA1* + *sdhA2*.

**Figure 4:**
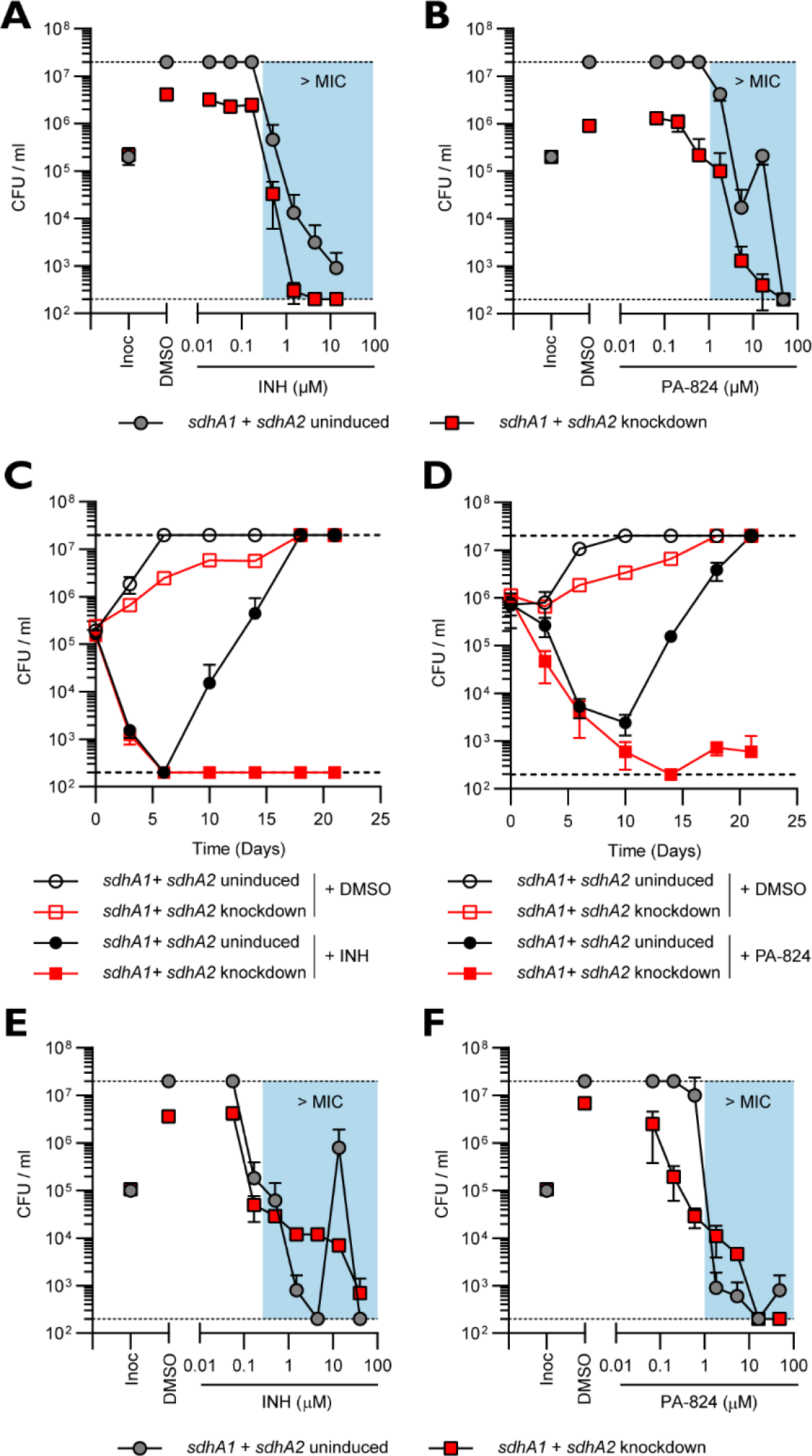
Impaired succinate oxidation by the joint transcriptional repression of *sdhA1* + *sdhA2* alters INH and PA-824 susceptibility and resistance in *M. tuberculosis*. (A and B) The effect of the transcriptional repression of *sdhA1* + *sdhA2* on the susceptibility of *M. tuberculosis* to INH **(A)** and PA-824 **(B)** when gene knockdown was induced simultaneously with antibiotic challenge. Cultures were grown in 96-well plates from a starting OD600 of 0.005 with 0 or 100 ng/ml ATc and a 7-point, 3-fold dilution gradient of each drug. Viability was determined after 10 days of incubation and CFU/ml were enumerated after 5 weeks. Results are mean ± SD of two replicates. **(C and D)** Viability of *M. tuberculosis sdhA1* + *sdhA2* knockdown strains treated with INH **(C)** or PA- 824 **(D)**. Cultures were grown in 7H9-OADC-PAN-KAN media in 10 ml volumes from a starting OD600 of 0.005. Knockdown was induced with 100 ng/ml ATc on day 0 and CFU/ml were determined on stated days. INH was used at 13.5 µM and PA-824 at 1.8 µM. Results are the mean and standard deviation of three replicates and are representative of three experiments. **(E and F)** Susceptibility of *M. tuberculosis* transcriptionally repressing *sdhA1* + *sdhA2* to INH **(E)** and PA-824 **(F)** when cells were pre-depleted of SDH enzymes prior to antibiotic challenge. Cultures were treated with 0 or 100 ng/ml ATc for 6 days to deplete cells of SDH enzymes, before harvesting cells and inoculating them into 96-well plates containing a 7-point, 3-fold dilution gradient of each drug and 0 or 100 ng/ml ATc at a starting OD600 of 0.005. Viability was determined by plating for CFU/ml after a further 10 days of incubation. Results are the means and standard deviations of two replicates. Blue boxes in **A**, **B**, **E** and **F** denote concentrations above the MIC of the no knockdown control. Dashed horizontal lines represent the upper and lower levels of detection. Inoc: CFU/ml at inoculation (i.e., time = 0).

To address this, we considered whether depleting cells of CRISPRi target enzymes prior to antibiotic challenge affected the interactions observed. We exploited the known synthetically lethality between the two terminal oxidases (the cytochrome *bc*1:*aa*3 complex and cytochrome *bd*) (13, 46, 62, 63) as a positive control. Killing of a *M. tuberculosis* strain expressing a sgRNA targeting cytochrome *bd* by the *bc*1 inhibitors, Q203 and TB47 (64), was only observed when cells were pre-depleted of cytochrome *bd*, and not when knockdown was induced simultaneously with antibiotic challenge (Figure S2). Therefore, we pre-depleted *M. tuberculosis* of SDH enzymes by inducing CRISPRi for 6 days before challenging cultures with the TB drugs listed above. Under these conditions, the dual depletion of Sdh1 + Sdh2 attenuated the bactericidal activity of INH against *M. tuberculosis* at concentrations above the MIC, while no difference in susceptibility was seen at concentrations below the MIC (Figure 4E). Note that the large ‘jump’ in CFU/ml for the no knockdown control at 13.5 µM INH was due to the emergence of INH resistance in one well. Similarly, depletion of Sdh1 + Sdh2 attenuated the bactericidal activity of PA-824 at concentrations above the MIC but, interestingly, resulted in increased killing at concentrations below the MIC of the knockdown control (Figure 4F). Moreover, the depletion of Sdh1 + Sdh2 attenuated the activity of all three other cell wall inhibitors tested: EMB, ETH, and SQ109 (Figure 5A-C). Combined, these data demonstrate that impaired succinate oxidation attenuates the bactericidal activity of cell wall inhibitors against *M. tuberculosis*.

**Figure 5:**
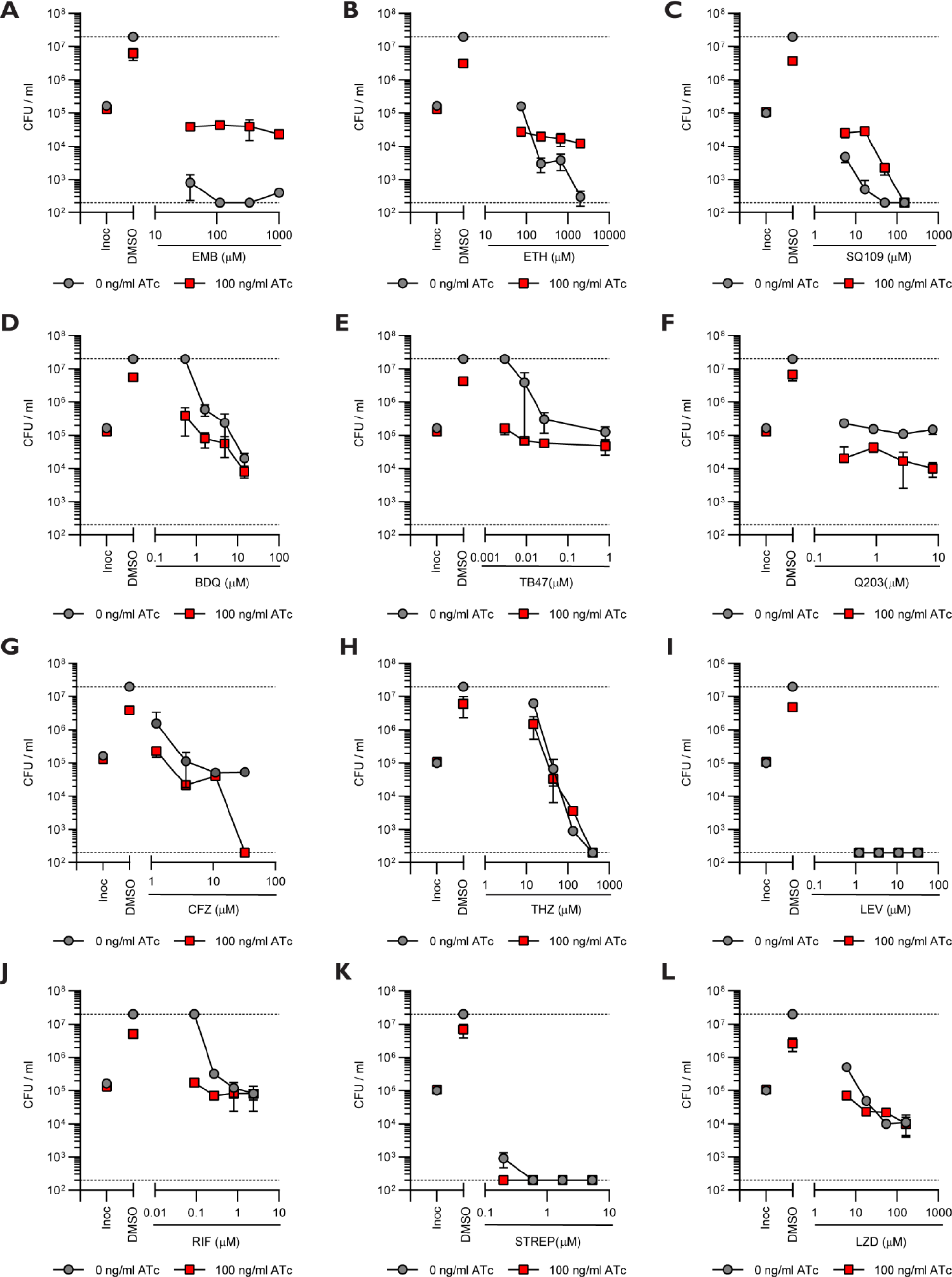
Impaired succinate oxidation by the dual depletion of Sdh1 + Sdh2 synergizes with bioenergetic inhibitors in *M. tuberculosis* but attenuates the activity of cell wall inhibitors. (A – C) Effect of the joint transcriptional repression of *sdhA1* + *sdhA2* on the susceptibility of *M. tuberculosis* to the cell wall inhibitors ethambutol (EMB) **(A)**, ethionamide (ETH) **(B)** and SQ109 **(C)**. **(D – H)** Effect of the joint transcriptional repression of *sdhA1* + *sdhA2* on the susceptibility of *M. tuberculosis* to bioenergetic inhibitors; BDQ **(D)**, TB47 **(E),** Q203 **(F)**, clofazimine (CFZ) **(G)** and thioridazine (THZ) **(H)**. **(I – L)** Effect of the joint transcriptional repression of *sdhA1* + *sdhA2* on the susceptibility of *M. tuberculosis* to inhibitors of DNA replication, transcription, or translation; levofloxacin (LEV) **(I)**, rifampicin (RIF) **(J)**, streptomycin (STREP) **(K)** and linezolid (LZD) **(L)**. Cultures were pre-depleted of SDH enzymes by inducing gene knockdown (0 or 100 ng/ml ATc) for 6 days and then inoculated into 96- well plates at a starting OD600 of 0.005 with 0 or 100 ng/ml ATc and a 7-point, 3-fold dilution gradient of each drug. Viability was determined after 10 days of incubation and CFU/ml were enumerated after 5 weeks. Dashed horizontal lines represent the upper and lower levels of detection. Results are the mean and standard deviation of two replicates. Inoc: CFU/ml at inoculation (i.e., time = 0).

In contrast, the dual depletion of Sdh1 + Sdh2 increased the susceptibility of *M. tuberculosis* to the bioenergetic inhibitors BDQ, TB47, Q203 and CFZ (Figure 5D-G). *M. tuberculosis* with both Sdh1 + Sdh2 depleted was hypersusceptible to growth inhibition by BDQ and TB47 at concentrations that were below the MIC of the no knockdown control (Figure 5D and E). Moreover, the depletion of Sdh1 + Sdh2 rendered *M. tuberculosis* susceptible to killing by Q203, resulting in a 0.5 – 1 log reduction in CFU/ml at concentrations that were bacteriostatic against the no knockdown control (Figure 5F). Increased killing of *M. tuberculosis* repressing *sdhA1* + *sdhA2* was also seen by CFZ at the highest concentration tested (32 µM) but not at lower concentrations (Figure 5G). Finally, impaired succinate oxidation did not alter the susceptibility of *M. tuberculosis* to the remainder of the other antimicrobials tested (Figure 5H-L), except for synergizing with RIF at sub-MIC levels (Figure 5J).

## Discussion

Succinate oxidation is a major focal point in both the central carbon metabolism and respiratory chain of *M. tuberculosis*; yet its essentiality remains poorly characterized. Here, we address this by reporting on the roles and essentialities of three different enzymes (Sdh1, Sdh2 and Frd) predicted to be capable of catalyzing the oxidation of succinate to fumarate in *M. tuberculosis*, and on the consequences of impaired succinate oxidation. We show that Sdh1 is the primary aerobic SDH utilized by *M. tuberculosis*, although its loss can be compensated for by Sdh2 but not Frd. Simultaneous repression of both Sdh1 and Sdh2 is required to impair succinate oxidation and prevent growth. Our results demonstrate that succinate oxidation, catalyzed by Sdh1 or Sdh2, is the major contributor of electrons to the respiratory chain and is required for the optimal growth of *M. tuberculosis* on both fermentable and non-fermentable carbon sources. Finally, we show that impaired succinate oxidation both positively and negatively affects the susceptibility of *M. tuberculosis* to a variety of anti-TB drugs.

Sdh1 has previously been proposed to be the primary aerobic SDH in *M. tuberculosis*, with its deletion resulting in a minor growth impairment in media containing succinate as the sole carbon and energy source (40). Likewise, the transcriptional repression of *sdhA1* impaired the growth and OCR of *M. tuberculosis* in succinate containing media, validating our CRISPRi approach to studying the essentiality of SDH and FRD enzymes in *M. tuberculosis*. Notably, the deletion of *sdh1* was reported to result in an increased rate of respiration in *M. tuberculosis* (40), which may appear to contradict our finding that the single *sdhA1* knockdown strain consumes oxygen more slowly in succinate containing media. However, the previously described increased respiration rate in the *Δsdh1* strain was only observed when oxygen levels fell below ∼40% dissolved oxygen, while oxygen consumption rates were similar at higher oxygen tensions (40). In our experiments using the same media (containing glycerol as an energy source), we observed no difference in OCR between the single *sdhA1* knockdown strain and the no knockdown control, thus reconciling the two findings.

In addition to validating Sdh1 is the primary aerobic SDH in *M. tuberculosis*, our work elucidates the compensatory mechanisms within succinate oxidation by demonstrating that Sdh2, but not Frd, can catalyze succinate oxidation in the absence of Sdh1. However, the growth and oxygen consumption impairments observed when Sdh2 is the only functional SDH enzyme present, suggest that Sdh2 is less efficient at catalyzing succinate oxidation than Sdh1. The molecular mechanisms underlying this are unclear, although both enzymes appear to use different reaction mechanisms to drive succinate oxidation (34–36). Moreover, these findings are in contrast to *M. smegmatis* where Sdh2 is proposed to have a higher affinity for succinate oxidation and is essential, while Sdh1 can be deleted with no identifiable phenotype (154). Finally, while our results suggest that Frd is unable to carry out succinate oxidation, additional biochemical evidence is needed to validate this.

While individually dispensable, the dual depletion of both Sdh1 and Sdh2 demonstrates the essentiality of succinate oxidation in *M. tuberculosis*. The double *sdhA1* + *sdhA2* knockdown strain was significantly impaired for growth across a range of carbon sources, including the fermentable carbon sources glycerol and glucose, as well as the non-fermentable carbon sources succinate and acetate. The improved growth on glucose and glycerol is likely due to substrate-level phosphorylation augmenting ATP levels. Importantly, mycobacterial growth and persistence *in vivo* is believed to be fueled by fatty acids (4, 39, 65), rather than glycerol and glycolytic carbon sources (66). Thus, the inability of *M. tuberculosis* to grow on acetate (a fatty acid surrogate (67)) when succinate oxidation is impaired, suggests that inhibition of SDH activity may prevent growth *in vivo*. This is supported by the previously reported survival defect of the *M. tuberculosis* Δ*sdh1* strain in mice lungs (40). However, this needs to be investigated further using the double *sdhA1* + *sdhA2* knockdown strain in *M. tuberculosis* infection models.

The simultaneous knockdown of *sdhA1* + *sdhA2* reduced the OCR of *M. tuberculosis* by approximately 70%, demonstrating that succinate oxidation is essential for the normal function of the respiratory chain. This is interesting as the ETC of *M. tuberculosis* is known for its significant plasticity, with the bacterium encoding multiple other primary dehydrogenases in addition to its two SDH enzymes (32, 68) that could theoretically compensate for reduced SDH activity. The simplest explanation for the observed reduction in OCR is that the majority of electrons are donated to the respiratory chain of *M. tuberculosis* through the oxidation of succinate to fumarate. However, NADH is generally considered to be the primary electron donor for aerobic growth in mycobacteria (68). Therefore, it is possible that the reduced OCR in the double *sdhA1* + *sdhA2* knockdown strain is reflective of an overall reduction in central carbon metabolism and respiration to circumvent the loss of SDH activity. Succinate oxidation appears to be an invariant component of mycobacterial central carbon metabolism with little option to re-route around (28) and we hypothesize that cells downregulate metabolism and respiration to prevent an accumulation of succinate, which may disrupt the metabolic homeostasis of replicating cells. Regardless, our data supports the previous proposal that SDH is the master regulator of respiration in *M. tuberculosis* (40).

The essentiality of succinate oxidation for the optimal growth and ETC activity of *M. tuberculosis* complements previous reports on the requirement for SDH / FRD enzymes for mycobacterial persistence (3, 33, 37, 40, 41, 67). While not investigated in this study due to technical limitations of using CRISPRi under hypoxia, several studies have reported the essentiality of succinate metabolism for the survival of *M. tuberculosis* under low oxygen tensions (3, 33, 37, 40, 41, 67). In particular, inhibition of SDH activity using the suicide inhibitor 3-nitropropionate (3-NP) resulted in time-dependent killing of *M. tuberculosis* during adaptation to hypoxia (67). In contrast, our results suggest that the mycobacterial SDH / FRD enzymes are dispensable for survival under nutrient starvation, despite Frd being strongly upregulated under this condition (38). However, a potential limitation of using transcriptional repression without concomitant protein degradation (69, 70) is that there may be residual SDH / FRD enzymes in the cell that could catalyze succinate oxidation. This is particularly relevant under non-replicating conditions (i.e., nutrient starvation) where the lower energetic requirement of cells (i.e., maintenance energy) (10, 11) means that even a small amount of residual enzyme may be sufficient to sustain persistence. Therefore, while our data argues against a role for SDH / FRD under nutrient starvation, this needs additional validation.

In addition to providing fundamental insights into the physiology of mycobacterial energy metabolism, the findings described in this study add to the complex narrative surrounding the interaction between modulating respiration and antimicrobial efficacy. Here, we showed that inhibition of succinate oxidation attenuated the activity of several cell wall inhibitors against *M. tuberculosis* including INH, ETH, EMB and SQ109. Notably, we did not observe attenuated killing for all bactericidal antibiotics tested (e.g., THZ, LZD) arguing against a general inhibition of killing by inducing bacteriostasis as has previously been reported (53). These results are consistent with previous findings that the inhibition of respiration by co-treating mycobacteria with respiratory inhibitors such as Q203 or BDQ attenuates the bactericidal activity of INH, EMB and ETH by preventing a drug-induced lethal ATP burst (55–57). Interestingly, other studies have demonstrated that inhibiting respiration through alternative mechanisms can potentiate INH killing in *M. tuberculosis* (14, 61), while both enhancing and inhibiting respiration can prevent the emergence of INH resistance (14, 60, 61). Despite this complexity, our results demonstrate a requirement for succinate oxidation in the susceptibility of *M. tuberculosis* to INH and other cell wall biosynthesis inhibitors under *in vitro* conditions. The reports that altered respiration influences the development of INH resistance in *M. tuberculosis* (14, 60, 61) support our observation that impaired succinate oxidation prevents the emergence of INH and PA-824 resistance, although this finding requires additional investigation.

Lastly, our results highlight novel chemical-genetic interactions between ETC components that could be exploited for drug development. *M. tuberculosis* with both Sdh1 + Sdh2 transcriptionally depleted was hypersusceptible to growth inhibition and / or killing by other bioenergetic inhibitors, namely BDQ, TB47, Q203 and CFZ, and to the frontline drug, RIF. The fact that these interactions were only observed when cells were pre-depleted of SDH enzymes demonstrates a previously unreported technical consideration for using CRISPRi to screen for chemical-genetic interactions in *M. tuberculosis*. Overall, these findings add to previous reports of synergistic interactions between ETC complexes (13, 58, 59, 63, 71–73) and support the proposal that targeting multiple components of mycobacterial energy generation could result in efficacious drug regimens (74).

In summary, our work demonstrates that succinate oxidation is required for the optimal growth of *M. tuberculosis* and further defines the roles and essentiality of SDH / FRD enzymes in mycobacteria. The simultaneous transcriptional repression of *sdhA1* + *sdhA2* significantly reduced the activity of the respiratory chain, had bacteriostatic consequences on cell viability, and influenced the susceptibility of *M. tuberculosis* to a variety of antibiotics. Combined, these findings provide useful insights into the physiology of energy metabolism of *M. tuberculosis* and its utility as a nascent area for antimicrobial development.

## Materials and Methods

### Bacterial strains and culture conditions

*Escherichia coli* strain MC1061 was used for the construction of all CRISPRi expression plasmids. *E. coli* MC1061 was grown at 37°C in liquid Luria-Bertani broth (LB) with agitation (200 rpm) or on solid Luria-Bertani agar (LBA) supplemented with 1.5% (w/v) agar. CRISPRi plasmids were selected for and maintained with 50 µg/ml kanamycin.

The *M. tuberculosis* H37Rv auxotrophic-attenuated derivative mc^2^6230 (Δ*RD1*, Δ*panCD*) (75) was grown in Middlebrook 7H9 broth containing OADC (0.005% oleic acid, 0.5% bovine serum albumin (BSA) (Sigma), 0.2% dextrose, 0.085% catalase) and 0.05% tyloxapol. Media was supplemented with 25 µg/ml pantothenic acid (PAN). For culturing *M. tuberculosis* mc^2^6230 on single carbon sources, a base of 7H9 liquid media with 0.5% BSA, 0.085% NaCl, 0.05% tyloxapol and 25 µg/ml PAN was supplemented with either glycerol (0.2 %), glucose (20 mM), succinate (30 mM), acetate (0.2%), or a combination of acetate (0.1%) and glucose (10 mM). Cultures were maintained in 10 ml volumes in 30 ml inkwells at 37°C with agitation (140 rpm). CRISPRi plasmids were maintained in *M. tuberculosis* with 25 µg/ml kanamycin (KAN). When required, gene knockdown was induced by adding anhydrotetracycline (ATc) at 100 ng/ml. Solid media was Middlebrook 7H11 agar + OADC-PAN-KAN. For CFU determination, cultures were 10-fold serially diluted along a four-point dilution curve in 7H9, and 5 µl of each dilution was spotted onto 7H11 agar. Colonies were counted after five weeks of incubation at 37°C.

### Antimicrobial compounds

Rifampicin (RIF), isoniazid (INH), streptomycin (STREP), clofazimine (CFZ), thioridazine (THZ), levofloxacin (LEV), pretomanid (PA-824), ethionamide (ETH), kanamycin (KAN), ATc and SQ109 were purchased from Sigma-Aldrich. Linezolid (LZD) and ethambutol (EMB) were obtained from Selleck Chemicals. Bedaquiline (BDQ) was obtained from Toronto Research Chemicals. TB47 was a gift from Tianyu Zhang (Guangzhou Institutes of Biomedicine and Health, Chinese Academy of Sciences, China). Q203 and ND-011992 were gifts from Kevin Pethe (Lee Kong Chian School of Medicine, Nanyang Technological University, Singapore). All antibiotic stocks were prepared in DMSO, except for STREP, THZ and KAN which were prepared in H2O. ATc was dissolved in 70% ethanol.

### Construction of CRISPRi knockdown plasmids and strains

For transcriptional knockdowns in *M. tuberculosis*, a 20-25 bp sequence downstream of a permissible PAM sequence was identified in the non-template strand of *sdhA1* (Rv0248c), *sdhA2* (Rv3318) and *frdA* (Rv1553). The 20 – 25 bp sequence from the template and non-template strand were ordered as oligos with 5’ GGGA and AAAC overhangs, respectively. Information for individual sgRNA (i.e., target sequence, PAM sequence, and predicted PAM strength) is provided in Table S1. Oligos used for plasmid construction are listed in Table S2. Oligos were annealed and cloned into CRISPRi plasmids (i.e., pJLR965) using BsmB1 Golden Gate cloning, sequence verified, and transformed into *M. tuberculosis* strain mc^2^6230 as previously described (45). Plasmids are listed in Table S3.

Multiplexed plasmids that express two or three sgRNA targeting a combination of *sdhA1*, *sdhA2*, and *frdA* were constructed by cloning additional sgRNAs into a Sap-1 based Golden Gate cloning site (43). Plasmids that express either a total of two or three sgRNA were constructed using specific sets of primers to ensure that overhangs generated following Sap1 digestion would facilitate plasmid reassembly in a specific sgRNA order. In brief, multiplex plasmids that express a total of two sgRNAs were constructed by amplifying the additional sgRNA module (transcriptional promoter, sgRNA scaffold and transcriptional terminator) from the appropriate CRISPRi plasmid using Phusion polymerase and the following primer pair (sgRNA-2 = MMO120 + MMO121) as described in Table S4. The amplified sgRNA module was purified and cloned into the CRISPRi plasmid that expresses the appropriate sgRNA partner using golden gate cloning with Sap1, as described in Table S5. Multiplexed plasmids that express a total of three sgRNAs were constructed by amplifying the additional sgRNA modules from the appropriate CRISPRi plasmids as above using the following primer pairs (sgRNA-2 = MMO120 + MMO123 and sgRNA-3 = MMO122 + MMO121). The amplified sgRNA modules were purified and cloned into the CRISPRi plasmid that expresses the appropriate sgRNA partner as above. Constructed plasmids were sequence verified and transformed into *M. tuberculosis* strain mc^2^6230 as previously described (45). All constructed plasmids are listed in Table S3.

### RNA Extraction and mRNA quantification by qPCR

*M. tuberculosis* single and multiplexed *sdhA1*, *sdhA2*, and *frdA* CRISPRi strains were inoculated at an OD600 of 0.1 in 10 ml 7H9-OADC media containing 100 ng/ml ATc. Cultures were harvested for RNA extraction after three days of gene knockdown as previously described (45). RNA was extracted using TRIzol reagent (Invitrogen) and purified using the RNA Clean and Concentrator kit (Zymo) as previously described (45). Samples were DNase treated using the TURBO DNA-free kit (Invitrogen) and confirmed to be DNA free by PCR using 1 µl of extracted RNA as a template with the primer combination of MMO200+MMO201.

cDNA was synthesized from the RNA using the SuperScript IV VILO Master Mix (Invitrogen) according to the manufacturer’s instructions. qPCR reactions were performed in 384 well plates in a ViiA7 Thermocycler using the Invitrogen PowerUp SYBR Green Master Mix as previously described (45). Primer sequences are listed in Table S2.2. All qPCR primer pairs and cDNA masses tested were experimentally validated to be within the linear range of the assay. Signals were normalized to the housekeeping gene *sigA* and quantified by the 2^ΔΔCt^ method.

### Nutrient Starvation Experiments

The survival of *M. tuberculosis* knockdown strains under conditions of nutrient starvation was determined using previously published protocols (38). Briefly, *M. tuberculosis* knockdown strains were inoculated at a starting OD600 of 0.005 in complete medium (7H9 + OADC-PAN-KAN) in the presence of 100 ng/ml ATc to induce CRISPRi and deplete cells of SDH and FRD enzymes. After 8 days, cultures were harvested and washed twice in PBS + 0.05% tyloxapol. Cells were resuspended in PBS-Tyloxapol and nutrient starved by inoculating into inkwells containing PBS-Tyloxapol, KAN, and 100 ng/ml ATc at an OD600 of 0.01 (i.e., approximately 10^5^ CFU/ml). PAN was not included as *M. tuberculosis* mc^2^6230 has previously been shown not to require PAN under conditions of nutrient starvation (76, 77). Cultures were incubated at 37°C without shaking for 8 weeks and viability measured by plating for CFUs every 7 days.

### Oxygen Consumption Rate Measurements

Oxygen consumption rate measurements of exponentially growing *M. tuberculosis* cells were performed as previously described using an Oroboros O2k FluoRespirometer (58, 64). In brief, *M. tuberculosis* strains expressing sgRNA targeting *sdhA1*, *sdhA2*, and *frdA* in single or multiplexed constructs were inoculated at a starting OD600 of 0.005 in 10 ml 7H9 media supplemented with either OADC, glycerol or succinate, and 0 or 100 ng/ml ATc. Cultures were grown for 8 days until the *sdhA1* + *sdhA2* double knockdown strain reached an OD600 of ∼0.2. All cultures were adjusted to an OD600 of 0.2 in supplemented 7H9 media on the day of experimentation from log-phase cultures. 2 ml of culture was added to each measurement chamber, which contains an O2 sensing clark-type electrode. Oxygen consumption rates were monitored for 15 minutes once oxygen consumption reached a steady state. All measurements were made at 37°C with 750 rpm in a closed chamber and a data recording interval of 2 s^-1^. The combined inhibition of both terminal oxidases with Q203 (400 nM) and ND-011992 (100 µM) was used to demonstrate complete inhibition of oxygen consumption. Chemicals were added after 10 minutes of respiration at a steady state through the injection port of the stoppers using Hamilton syringes. Oxygen consumption rates were calculated using the Oroboros Data Lab software.

### Antibiotic susceptibility testing

The Minimum Inhibitory Concentrations (MIC) and Minimum Bactericidal Concentrations (MBC) of the *M. tuberculosis sdhA1* + *sdhA2* CRISPRi strain was determined as previously described (45). In brief, *M. tuberculosis* was inoculated into a 96-well plate containing 7H9 + OADC-PAN-KAN media at an OD600 of 0.005 in a final volume of 150 µl with 0 or 100 ng/ml ATc. Antibiotics were dispensed from a 9-point, 3-fold dilution gradient into each well, with a maximum of 2% DMSO. Cultures were incubated at 37 °C for 10 days without agitation. On day 10, OD600 values were measured in a Varioskan LUX microplate reader and MIC values for the no knockdown controls were calculated using the Gompertz equation (78). For MBC determination, culture was removed from rows containing 0 and 100 ng/ml ATc at day 10, 10-fold serially in 7H9 and 5 µL of each dilution spotted onto 7H11 agar. Colonies were counted after five weeks of incubation at 37 °C.

For pre-knockdown experiments, the *M. tuberculosis sdhA1* + *sdhA2* CRISPRi strain was grown in 7H9-OADC-PAN-KAN media in 10 ml volumes from a starting OD600 of 0.005 for 6 days prior to antibiotic challenge. Cultures were treated with 0 or 100 ng/ml ATc on day 0 to induce gene silencing. After 6 days, cultures were inoculated at an OD600 of 0.005 into a 96-well plate containing 7H9 + OADC-PAN-KAN media, 0 or 100 ng/ml ATc and 9-point, 3-fold dilution gradient of the respective antibiotic. Plates were incubated for 10 days before plating for CFU/ml as described above.

### Time-kill experiments

*M. tuberculosis* cultures were inoculated into inkwells containing 10 ml 7H9 media supplemented with OADC-PAN-KAN at a starting OD600 of 0.005. Cultures were treated with 0 or 100 ng/ml ATc and with selected antibiotics at the specified concentrations. Bacterial viability was determined by plating onto 7H11 agar plates at specified time points and enumerating CFU after 5 weeks of incubation at 37°C. For pre-knockdown experiments, strains were incubated in the presence of 0 or 100 ng/ml ATc for 6 days as described above before being inoculated into inkwells.

## Acknowledgements

This work was financially supported by the Maurice Wilkins Centre for Molecular Biodiscovery, the Marsden Fund (Royal Society of New Zealand) and the Health Research Council of New Zealand. C.A was supported by a University of Otago Doctoral Scholarship and a William Georgetti Scholarship.

We have no conflict of interests to declare.

## Supplementary Information

**Figure S1:**
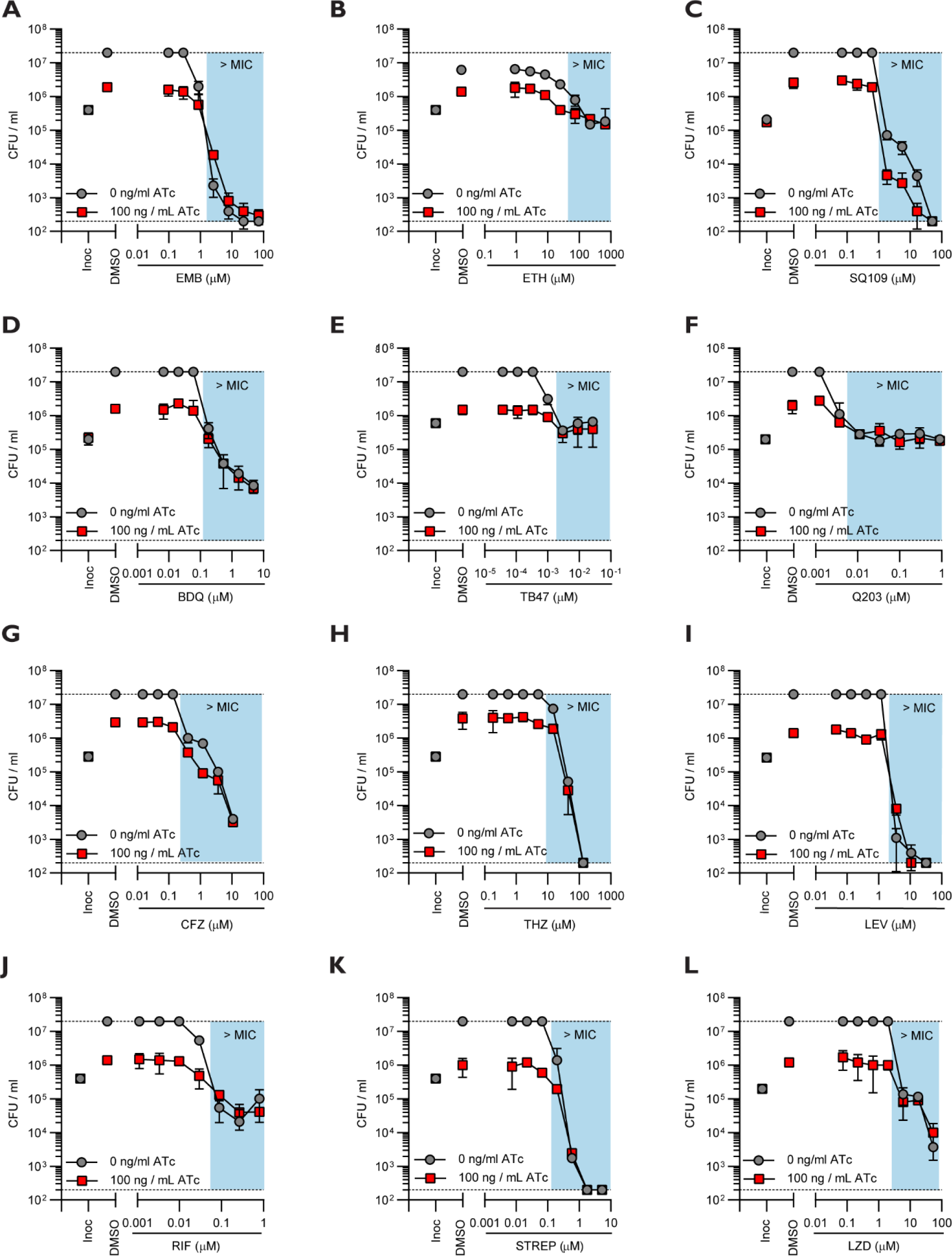
Impact of impaired succinate oxidation on the activity of various TB drugs against *M. tuberculosis*. Effect of the simultaneous transcriptional repression of *sdhA1* + *sdhA2* on the susceptibility of *M. tuberculosis* to various TB drugs. Cultures were grown in 96-well plates from a starting OD600 of 0.005 with a 7-point, 3-fold dilution gradient of each drug. Gene knockdown was indued on day 0 with 0 or 100 ng/ml ATc. Viability was determined after 10 days of incubation. Blue boxes denote concentrations above the MIC of the no knockdown control. Dashed horizontal lines represent the upper and lower levels of detection. Results are the mean and standard deviation of two replicates. EMB; ethambutol, ETH; ethionamide, BDQ; bedaquiline, CFZ; clofazimine, THZ; thioridazine, LEV; levofloxacin, RIF; rifampicin, STREP; streptomycin, LZD; linezolid. Inoc: CFU/ml at inoculation (i.e., time = 0).

**Figure S2:**
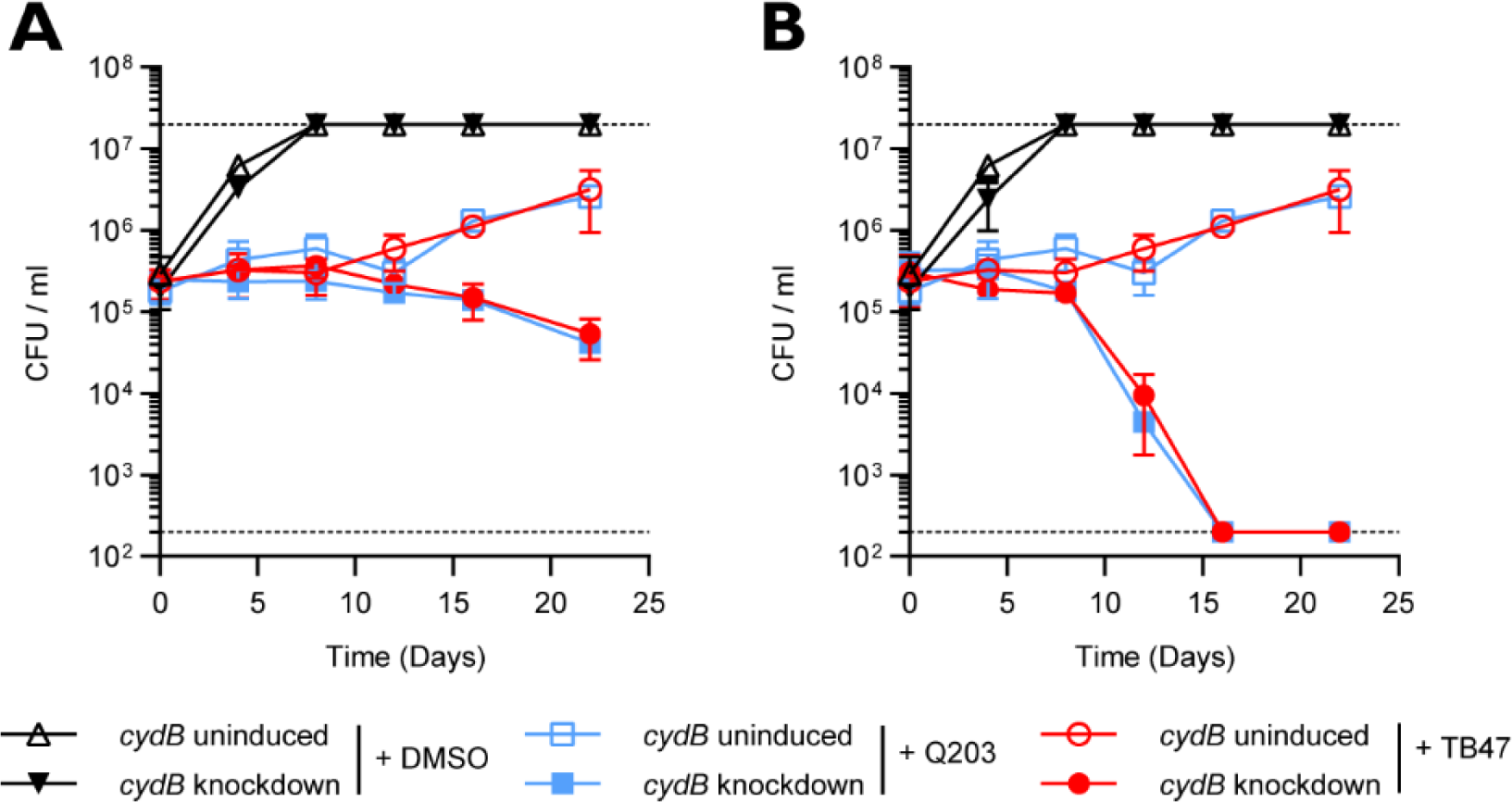
Impact of pre-depleting cells of CRISPRi target enzymes before antibiotic challenge. **(A)** Susceptibility of *M. tuberculosis* transcriptionally repressing cytochrome *bd* to Q203 (400 nM) and TB47 (400 nM) when knockdown was induced with 100 ng/ml ATc at day 0. **(B)** Susceptibility of *M. tuberculosis* transcriptionally repressing cytochrome *bd* to Q203 (400 nM) and TB47 (400 nM) when cells were pre-depleted of cytochrome *bd* by inducing gene knockdown (100 ng/ml ATc) for 6 days prior to antibiotic challenge. Cultures were grown in 7H9-OADC-PAN-KAN media in 10 ml volumes from a starting OD600 of 0.005. CFU/ml were determined on stated days and enumerated after 3 weeks of incubation. Dashed horizontal lines represent the upper and lower limits of detection. Results are the means and standard deviations of two replicates.

**Table S1:**
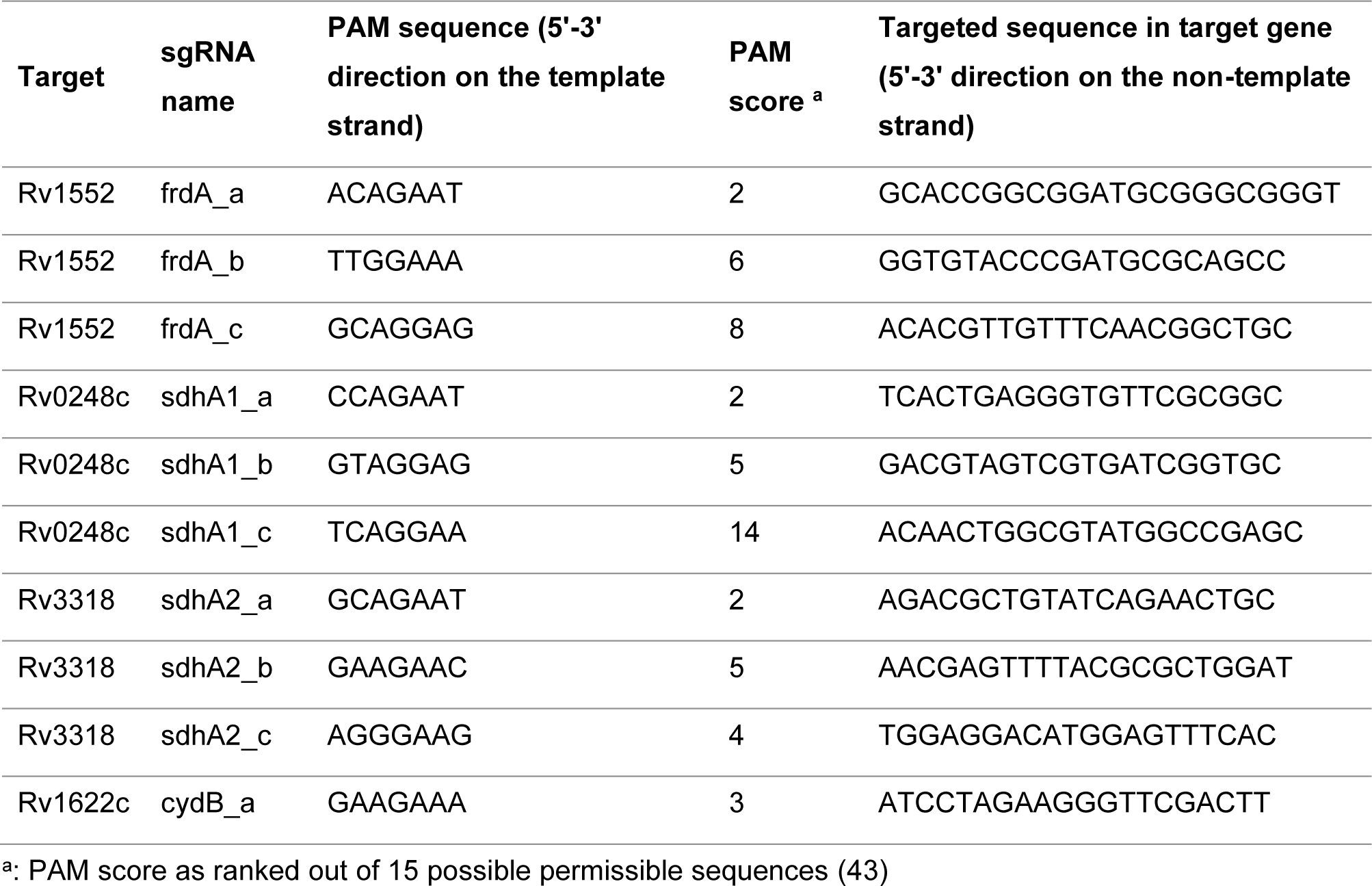
sgRNA targeting sdhA1, sdhA2, frdA and cydB in M. tuberculosis

**Table S2:**
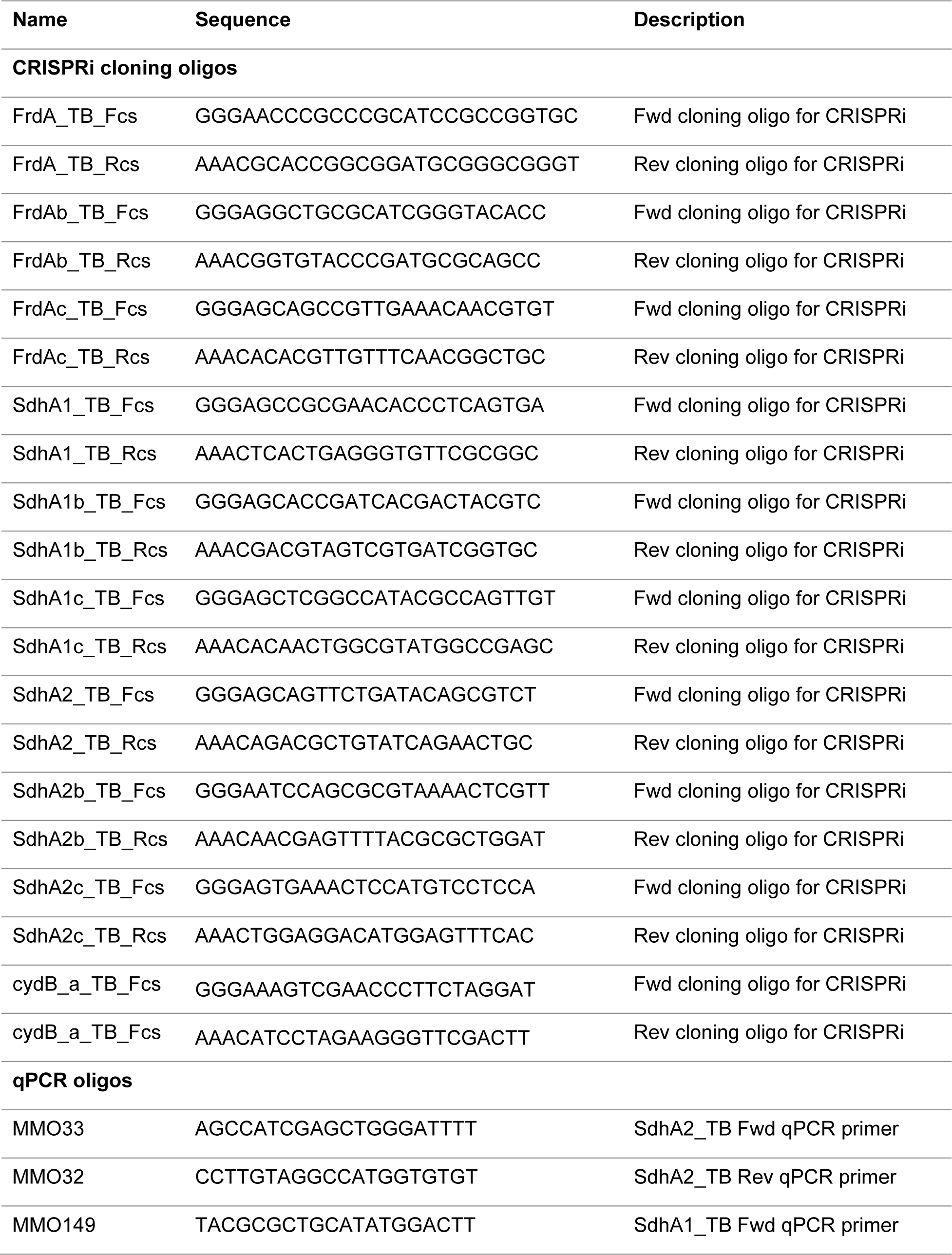
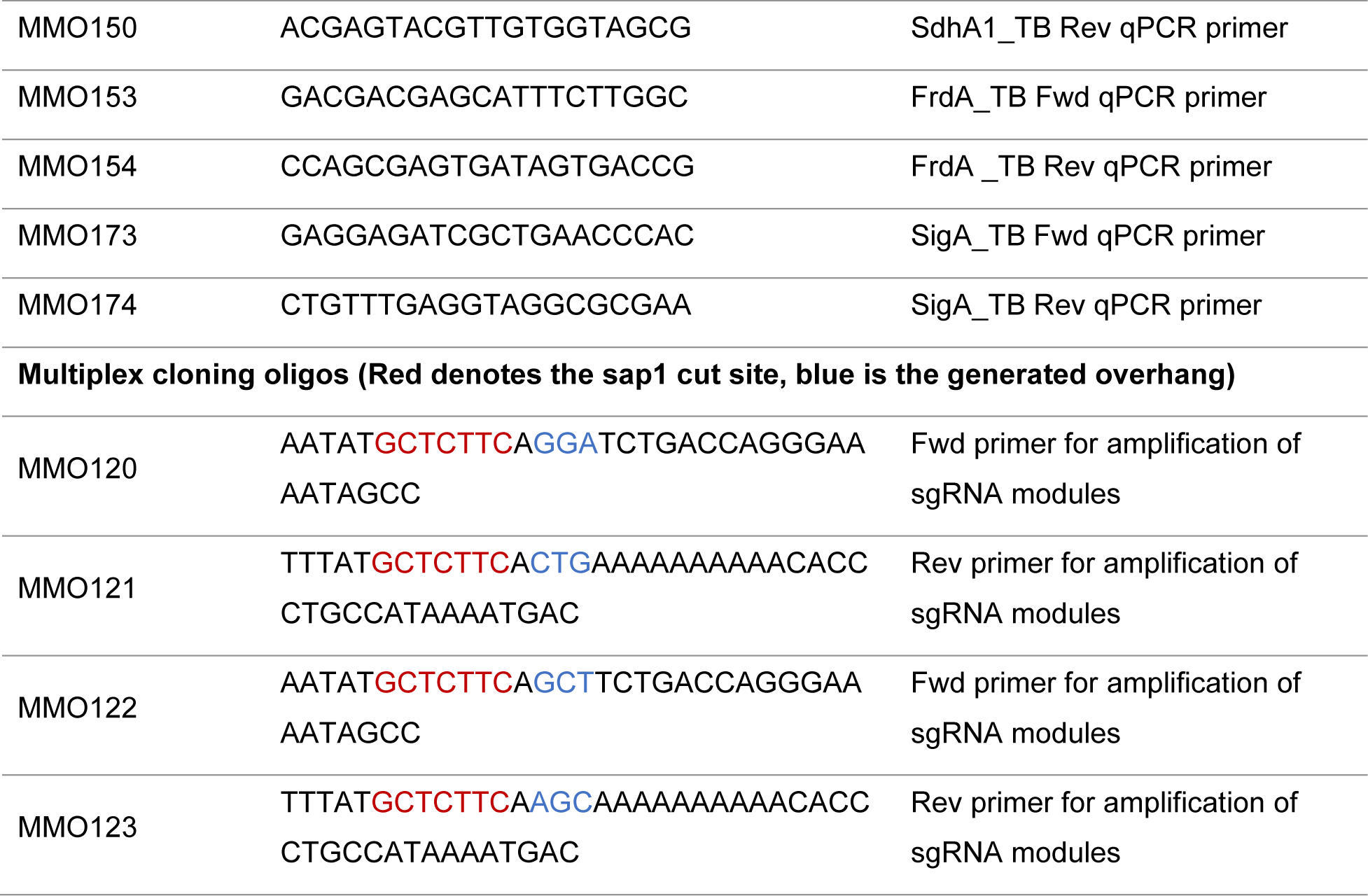
Oligos used in this study

**Table S3:** Plasmids used in this study

| Name | Description | Plasmid Backbone | Reference |
| --- | --- | --- | --- |
| pJLR965 | CRISPRi cloning vector for *M. tuberculosis*- vector expressing dCas9 and non-targeting sgRNA |  | (43) |
| **Single sgRNA plasmids** |  |  |  |
| pCi7 | CRISPRi vector expressing dCas9 + sgRNA targeting *mmpL3* | pJLR965 | (45) |
| pCi5 | CRISPRi vector expressing dCas9 + sgRNA (*frdA_a*) targeting *frdA* | pJLR965 | This study |
| pCi8 | CRISPRi vector expressing dCas9 + sgRNA (*sdhA1_a*) targeting *sdhA1* | pJLR965 | This study |
| pCi9 | CRISPRi vector expressing dCas9 + sgRNA (*sdhA2_a*) targeting *sdhA2* | pJLR965 | This study |
| pCi23 | CRISPRi vector expressing dCas9 + sgRNA (*frdA_b*) targeting *frdA* | pJLR965 | This study |
| pCi24 | CRISPRi vector expressing dCas9 + sgRNA (*frdA_b*) targeting *frdA* | pJLR965 | This study |
| pCi25 | CRISPRi vector expressing dCas9 + sgRNA (*sdhA1_b*) targeting *sdhA1* | pJLR965 | This study |
| pCi26 | CRISPRi vector expressing dCas9 + sgRNA (*sdhA1_c*) targeting *sdhA1* | pJLR965 | This study |
| pCi27 | CRISPRi vector expressing dCas9 + sgRNA (*sdhA2_b*) targeting *sdhA2* | pJLR965 | This study |
| pCi28 | CRISPRi vector expressing dCas9 + sgRNA (*sdhA2_c*) targeting *sdhA2* | pJLR965 | This study |
| pCi95 | CRISPRi vector expressing dCas9 + sgRNA (*cydB*) targeting *cydB* | pJLR965 | This study |
| **Multiplexed plasmids** |  |  |  |
| pCiMX15 | CRISPRi vector expressing dCas9 + sgRNAs (*sdhA1_a* and *sdhA2_a*) | pCi8 | This study |
| pCiMX22 | CRISPRi vector expressing dCas9 + sgRNAs (*frdA_b* and *sdhA1_a*) | pCi23 | This study |
| pCiMX23 | CRISPRi vector expressing dCas9 + sgRNAs (*frdA_b* and *sdhA2_a*) | pCi23 | This study |
| pCiMX24 | CRISPRi vector expressing dCas9 + sgRNAs (*frdA_b*, *sdhA1_a* and *sdhA2_*) | pCi23 | This study |

**Table S4:** Amplification of target sgRNA for multiplexed cloning

| Component | 50 µl RxN (µl) |  |  |
| --- | --- | --- | --- |
| H<sub>2</sub>O | 31 |  |  |
| 5x Phusion GC Buffer | 10 |  |  |
| 10 mM dNTPs | 1 |  |  |
| Fwd Primer (10 µM stock) | 2.5 |  |  |
| Rev Primer (10 µM stock) | 2.5 |  |  |
| pCi plasmid with target sgRNA | 1 µl |  |  |
| DMSO | 1.5 |  |  |
| Phusion Poln | 0.5 |  |  |
| Total | 50 |  |  |
| Thermocycler instructions |  |  |  |
| Step | Temp | Time | Cycles |
| Initial Denaturation | 98 °C | 30 sec | 1 |
| Denaturation | 98 °C | 10 sec | 35 |
| Annealing | 60 °C | 30 sec |  |
| Extension | 72 °C | 1 min |  |
| Final Extension | 72 °C | 10 min | 1 |
| Hold | 4 °C | hold |  |

**Table S5:**
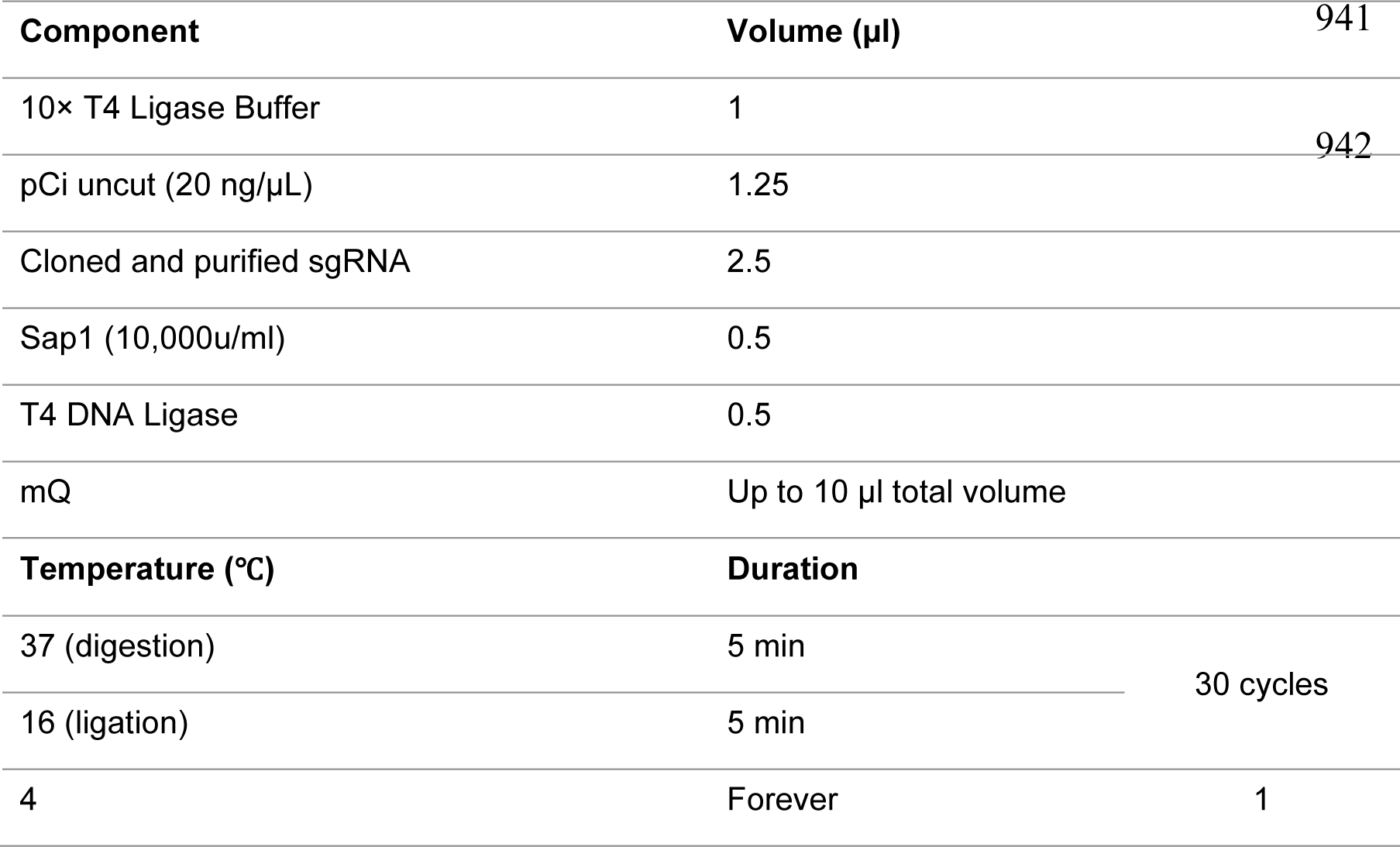
Multiplex golden gate cloning

## References

1. World Health Organisation. 2020. Global tuberculosis report.

2. World Health Organization. 2010. Guidelines for the treatment of tuberculosis: Fourth Edition

3. Rittershaus ESC, Baek S-H, Krieger IV, Nelson SJ, Cheng Y-S, Nambi S, Baker RE, Leszyk JD, Shaffer SA, Sacchettini JC, Sassetti CM. 2018. A Lysine Acetyltransferase Contributes to the Metabolic Adaptation to Hypoxia in *Mycobacterium tuberculosis*. Cell Chemical Biology 25:1495–1505.e3.

4. Munoz-Elias EJ, McKinney JD. 2005. *Mycobacterium tuberculosis* isocitrate lyases 1 and 2 are jointly required for in vivo growth and virulence. Nat Med 11:638–44.

5. Marrero J, Rhee KY, Schnappinger D, Pethe K, Ehrt S. 2010. Gluconeogenic carbon flow of tricarboxylic acid cycle intermediates is critical for *Mycobacterium tuberculosis* to establish and maintain infection. Proc Natl Acad Sci U S A 107:9819–24.

6. Ruecker N, Jansen R, Trujillo C, Puckett S, Jayachandran P, Piroli GG, Frizzell N, Molina H, Rhee KY, Ehrt S. 2017. Fumarase Deficiency Causes Protein and Metabolite Succination and Intoxicates *Mycobacterium tuberculosis*. Cell Chem Biol 24:306–315.

7. McKinney JD, Honer zu Bentrup K, Munoz-Elias EJ, Miczak A, Chen B, Chan W, Swenson D, Sacchettini JC, Jacobs Jr. WR, Russell DG. 2000. Persistence of *Mycobacterium tuberculosis* in macrophages and mice requires the glyoxylate shunt enzyme isocitrate lyase. Nature 406:735–738.

8. Munoz-Elias EJ, Upton AM, Cherian J, McKinney JD. 2006. Role of the methylcitrate cycle in *Mycobacterium tuberculosis* metabolism, intracellular growth, and virulence. Mol Microbiol 60:1109–22.

9. Venugopal A, Bryk R, Shi S, Rhee K, Rath P, Schnappinger D, Ehrt S, Nathan C. 2011. Virulence of *Mycobacterium tuberculosis* depends on lipoamide dehydrogenase, a member of three multienzyme complexes. Cell Host Microbe 9:21–31.

10. Gengenbacher M, Rao SP, Pethe K, Dick T. 2010. Nutrient-starved, non- replicating *Mycobacterium tuberculosis* requires respiration, ATP synthase and isocitrate lyase for maintenance of ATP homeostasis and viability. Microbiology 156:81–7.

11. Rao SP, Alonso S, Rand L, Dick T, Pethe K. 2008. The protonmotive force is required for maintaining ATP homeostasis and viability of hypoxic nonreplicating *Mycobacterium tuberculosis*. Proc Natl Acad Sci U S A 105:11945–11950.

12. Koul A, Vranckx L, Dendouga N, Balemans W, Van den Wyngaert I, Vergauwen K, Gohlmann HW, Willebrords R, Poncelet A, Guillemont J, Bald D, Andries K. 2008. Diarylquinolines are bactericidal for dormant mycobacteria as a result of disturbed ATP homeostasis. J Biol Chem 283:25273–80.

13. Kalia NP, Hasenoehrl EJ, Ab Rahman NB, Koh VH, Ang MLT, Sajorda DR, Hards K, Gruber G, Alonso S, Cook GM, Berney M, Pethe K. 2017. Exploiting the synthetic lethality between terminal respiratory oxidases to kill *Mycobacterium tuberculosis* and clear host infection. Proc Natl Acad Sci U S A 114:7426–7431.

14. Sukheja P, Kumar P, Mittal N, Li SG, Singleton E, Russo R, Perryman AL, Shrestha R, Awasthi D, Husain S, Soteropoulos P, Brukh R, Connell N, Freundlich JS, Alland D. 2017. A Novel Small-Molecule Inhibitor of the *Mycobacterium tuberculosis* Demethylmenaquinone Methyltransferase MenG Is Bactericidal to Both Growing and Nutritionally Deprived Persister Cells. mBio 8.

15. Andries K, Verhasselt P, Guillemont J, Gohlmann HW, Neefs JM, Winkler H, Van Gestel J, Timmerman P, Zhu M, Lee E, Williams P, de Chaffoy D, Huitric E, Hoffner S, Cambau E, Truffot-Pernot C, Lounis N, Jarlier V. 2005. A Diarylquinonline Drug Active on the ATP Synthase of *Mycobacterium tuberculosis*. Science 307:223–227.

16. Koul A, Dendouga N, Vergauwen K, Molenberghs B, Vranckx L, Willebrords R, Ristic Z, Lill H, Dorange I, Guillemont J, Bald D, Andries K. 2007. Diarylquinolines target subunit c of mycobacterial ATP synthase. Nat Chem Biol 3:323–4.

17. Priess L, Langer JD, Yildiz O, Eckhardt-Strelau L, Guillemont JEG, Koul A, Meier T. 2015. Structure of the mycobacterial ATP Synthase F0 rotor ring in complex with the anti-TB drug bedaquiline. Sci Adv 1.

18. Manjunatha U, Boshoff HI, Barry CE. 2009. The mechanism of action of PA-824: Novel insights from transcriptional profiling. Commun Integr Biol 2:215–8.

19. Singh R, Manjunatha U, Boshoff HI, Ha YH, Niyomrattanakit P, Ledwidge R, Dowd CS, Lee IY, Kim P, Zhang L, Kang S, Keller TH, Jiricek J, Barry CE, 3rd. 2008. PA- 824 kills nonreplicating Mycobacterium tuberculosis by intracellular NO release. Science 322:1392–5.

20. Van den Bossche A, Varet H, Sury A, Sismeiro O, Legendre R, Coppee JY, Mathys V, Ceyssens PJ. 2019. Transcriptional profiling of a laboratory and clinical *Mycobacterium tuberculosis* strain suggests respiratory poisoning upon exposure to delamanid. Tuberculosis (Edinb) 117:18–23.

21. Stover CK, Warrener P, VanDevanter DR, Sherman DR, Arain TM, Langhorne MH, Anderson SW, Towell JA, Ying Yuan, McMurray DN, Kreiswirth BN, Barry CE, Baker WR. 2000. A small-molecule nitroimidazopyran drug candidate for the treatment of tuberculosis. Nature 405.

22. Matsumoto M, Hashizume H, Tomishige T, Kawasaki M, Tsubouchi H, Sasaki H, Shimokawa Y, Komatsu M. 2006. OPC-67683, a nitro-dihydro-imidazooxazole derivative with promising action against tuberculosis in vitro and in mice. PLoS Med 3:e466.

23. Diacon AH, Donald PR, Pym A, Grobusch M, Patientia RF, Mahanyele R, Bantubani N, Narasimooloo R, De Marez T, van Heeswijk R, Lounis N, Meyvisch P, Andries K, McNeeley DF. 2012. Randomized pilot trial of eight weeks of bedaquiline (TMC207) treatment for multidrug-resistant tuberculosis: long-term outcome, tolerability, and effect on emergence of drug resistance. Antimicrob Agents Chemother 56:3271–6.

24. Keam SJ. 2019. Pretomanid: First Approval. Drugs 79:1797–1803.

25. Ryan NJ, Lo JH. 2014. Delamanid: first global approval. Drugs 74:1041–5.

26. Jones D. 2013. Tuberculosis success. Nat Rev Drug Discov 12:175–6.

27. Conradie F, Diacon AH, Ngubane N, Howell P, Everitt D, Crook AM, Mendel CM, Egizi E, Moreira J, Timm J, McHugh TD, Wills GH, Bateson A, Hunt R, Van Niekerk C, Li M, Olugbosi M, Spigelman M, Nix TBTT. 2020. Treatment of Highly Drug-Resistant Pulmonary Tuberculosis. N Engl J Med 382:893–902.

28. Hards K, Adolph C, Harold LK, McNeil MB, Cheung CY, Jinich A, Rhee KY, Cook GM. 2020. Two for the price of one: Attacking the energetic-metabolic hub of mycobacteria to produce new chemotherapeutic agents. Prog Biophys Mol Biol doi:10.1016/j.pbiomolbio.2019.11.003.

29. Maklashina E, Cecchini G, Dikanov SA. 2013. Defining a direction: electron transfer and catalysis in *Escherichia coli* complex II enzymes. Biochim Biophys Acta 1827:668–78.

30. Lancaster CRD. 2013. The di-heme family of respiratory complex II enzymes. Biochim Biophys Acta 1827:679–87.

31. Maklashina E, Berthold DA, Cecchini G. 1998. Anaerobic expression of *Escherichia coli* succinate dehydrogenase: functional replacement of fumarate reductase in the respiratory chain during anaerobic growth. J Bacteriol 180:5989–5996.

32. Cook GM, Hards K, Vilcheze C, Hartman T, Berney M. 2014. Energetics of Respiration and Oxidative Phosphorylation in Mycobacteria. Microbiol Spectr 2.

33. Pecsi I, Hards K, Ekanayaka N, Berney M, Hartman T, Jacobs WR, Jr., Cook GM. 2014. Essentiality of succinate dehydrogenase in *Mycobacterium smegmatis* and its role in the generation of the membrane potential under hypoxia. MBio 5.

34. Hards K, Rodriguez SM, Cairns C, Cook GM. 2019. Alternate quinone coupling in a new class of succinate dehydrogenase may potentiate mycobacterial respiratory control. FEBS Lett 593:475–486.

35. Gong H, Gao Y, Zhou X, Xiao Y, Wang W, Tang Y, Zhou S, Zhang Y, Ji W, Yu L, Tian C, Lam SM, Shui G, Guddat LW, Wong LL, Wang Q, Rao Z. 2020. Cryo-EM structure of trimeric *Mycobacterium smegmatis* succinate dehydrogenase with a membrane-anchor SdhF. Nat Commun 11:4245.

36. Zhou X, Gao Y, Wang W, Yang X, Yang X, Liu F, Tang Y, Lam SM, Shui G, Yu L, Tian C, Guddat LW, Wang Q, Rao Z, Gong H. 2021. Architecture of the mycobacterial succinate dehydrogenase with a membrane-embedded Rieske FeS cluster. Proceedings of the National Academy of Sciences 118.

37. Watanabe S, Zimmermann M, Goodwin MB, Sauer U, Barry CE, 3rd, Boshoff HI. 2011. Fumarate reductase activity maintains an energized membrane in anaerobic Mycobacterium tuberculosis. PLoS Pathog 7:e1002287.

38. Betts J, Lukey, PT, Robb, LC, McAdam, RA, Duncan K. 2002. Evaluation of a nutrient starvation model of *Mycobacterium tuberculosis* persistence by gene and protein expression profiling. Mol Microbiol 43:717–731.

39. Schnappinger D, Ehrt S, Voskuil MI, Liu Y, Mangan JA, Monahan IM, Dolganov G, Efron B, Butcher PD, Nathan C, Schoolnik GK. 2003. Transcriptional Adaptation of *Mycobacterium tuberculosis* within Macrophages: Insights into the Phagosomal Environment. J Exp Med 198:693–704.

40. Hartman T, Weinrick B, Vilcheze C, Berney M, Tufariello J, Cook GM, Jacobs WR, Jr. 2014. Succinate dehydrogenase is the regulator of respiration in *Mycobacterium tuberculosis*. PLoS Pathog 10:e1004510.

41. Baek SH, Li AH, Sassetti CM. 2011. Metabolic regulation of mycobacterial growth and antibiotic sensitivity. PLoS Biol 9:e1001065.

42. Griffin JE, Gawronski JD, Dejesus MA, Ioerger TR, Akerley BJ, Sassetti CM. 2011. High-resolution phenotypic profiling defines genes essential for mycobacterial growth and cholesterol catabolism. PLoS Pathog 7:e1002251.

43. Rock JM, Hopkins FF, Chavez A, Diallo M, Chase MR, Gerrick ER, Pritchard JR, Church GM, Rubin EJ, Sassetti CM, Schnappinger D, Fortune SM. 2017. Programmable transcriptional repression in mycobacteria using an orthogonal CRISPR interference platform. Nat Microbiol 2:16274.

44. McNeil MB, Ryburn HWK, Harold LK, Tirados JF, Cook GM. 2020. Transcriptional inhibition of the F1Fo-type ATP synthase has bactericidal consequences on the viability of mycobacteria. Antimicrob Agents Chemother doi:10.1128/AAC.00492-20.

45. McNeil MB, Cook GM. 2019. Utilization of CRISPR Interference To Validate MmpL3 as a Drug Target in *Mycobacterium tuberculosis*. Antimicrob Agents Chemother 63.

46. Lee BS, Hards K, Engelhart CA, Hasenoehrl EJ, Kalia NP, Mackenzie JS, Sviriaeva E, Chong SMS, Manimekalai MSS, Koh VH, Chan J, Xu J, Alonso S, Miller MJ, Steyn AJC, Gruber G, Schnappinger D, Berney M, Cook GM, Moraski GC, Pethe K. 2020. Dual inhibition of the terminal oxidases eradicates antibiotic-tolerant *Mycobacterium tuberculosis*. EMBO Mol Med doi:10.15252/emmm.202013207:e13207.

47. Pethe K, Bifani P, Jang J, Kang S, Park S, Ahn S, Jiricek J, Jung J, Jeon HK, Cechetto J, Christophe T, Lee H, Kempf M, Jackson M, Lenaerts AJ, Pham H, Jones V, Seo MJ, Kim YM, Seo M, Seo JJ, Park D, Ko Y, Choi I, Kim R, Kim SY, Lim S, Yim SA, Nam J, Kang H, Kwon H, Oh CT, Cho Y, Jang Y, Kim J, Chua A, Tan BH, Nanjundappa MB, Rao SP, Barnes WS, Wintjens R, Walker JR, Alonso S, Lee S, Kim J, Oh S, Oh T, Nehrbass U, Han SJ, No Z, et al. 2013. Discovery of Q203, a potent clinical candidate for the treatment of tuberculosis. Nat Med 19:1157–60.

48. Kohanski MA, Dwyer DJ, Hayete B, Lawrence CA, Collins JJ. 2007. A common mechanism of cellular death induced by bactericidal antibiotics. Cell 130:797–810.

49. Dwyer DJ, Kohanski MA, Hayete B, Collins JJ. 2007. Gyrase inhibitors induce an oxidative damage cellular death pathway in *Escherichia coli*. Mol Syst Biol 3:91.

50. Kohanski MA, Dwyer DJ, Wierzbowski J, Cottarel G, Collins JJ. 2008. Mistranslation of membrane proteins and two-component system activation trigger antibiotic-mediated cell death. Cell 135:679–90.

51. Belenky P, Ye JD, Porter CB, Cohen NR, Lobritz MA, Ferrante T, Jain S, Korry BJ, Schwarz EG, Walker GC, Collins JJ. 2015. Bactericidal Antibiotics Induce Toxic Metabolic Perturbations that Lead to Cellular Damage. Cell Rep 13:968–80.

52. Dwyer DJ, Belenky PA, Yang JH, MacDonald IC, Martell JD, Takahashi N, Chan CT, Lobritz MA, Braff D, Schwarz EG, Ye JD, Pati M, Vercruysse M, Ralifo PS, Allison KR, Khalil AS, Ting AY, Walker GC, Collins JJ. 2014. Antibiotics induce redox-related physiological alterations as part of their lethality. Proc Natl Acad Sci U S A 111:E2100–9.

53. Lobritz MA, Belenky P, Porter CB, Gutierrez A, Yang JH, Schwarz EG, Dwyer DJ, Khalil AS, Collins JJ. 2015. Antibiotic efficacy is linked to bacterial cellular respiration. Proc Natl Acad Sci U S A 112:8173–80.

54. Feng X, Zhu W, Schurig-Briccio LA, Lindert S, Shoen C, Hitchings R, Li J, Wang Y, Baig N, Zhou T, Kim BK, Crick DC, Cynamon M, McCammon JA, Gennis RB, Oldfield E. 2015. Antiinfectives targeting enzymes and the proton motive force. Proc Natl Acad Sci U S A 112:E7073–82.

55. Lee BS, Kalia NP, Jin XEF, Hasenoehrl EJ, Berney M, Pethe K. 2019. Inhibitors of energy metabolism interfere with antibiotic-induced death in mycobacteria. Journal of Biological Chemistry 294:1936–1943.

56. Shetty A, Dick T. 2018. Mycobacterial Cell Wall Synthesis Inhibitors Cause Lethal ATP Burst. Front Microbiol 9:1898.

57. Zeng S, Soetaert K, Ravon F, Vandeput M, Bald D, Kauffmann JM, Mathys V, Wattiez R, Fontaine V. 2019. Isoniazid Bactericidal Activity Involves Electron Transport Chain Perturbation. Antimicrob Agents Chemother 63.

58. Hards K, Robson JR, Berney M, Shaw L, Bald D, Koul A, Andries K, Cook GM. 2015. Bactericidal mode of action of bedaquiline. J Antimicrob Chemother 70:2028–37.

59. Berney M, Hartman TE, Jacobs WR, Jr. 2014. A *Mycobacterium tuberculosis* cytochrome *bd* oxidase mutant is hypersensitive to bedaquiline. mBio 5:e01275–14.

60. Vilcheze C, Hartman T, Weinrick B, Jain P, Weisbrod TR, Leung LW, Freundlich JS, Jacobs WR, Jr. 2017. Enhanced respiration prevents drug tolerance and drug resistance in *Mycobacterium tuberculosis*. Proc Natl Acad Sci U S A 114:4495–4500.

61. Flentie K, Harrison GA, Tukenmez H, Livny J, Good JAD, Sarkar S, Zhu DX, Kinsella RL, Weiss LA, Solomon SD, Schene ME, Hansen MR, Cairns AG, Kulen M, Wixe T, Lindgren AEG, Chorell E, Bengtsson C, Krishnan KS, Hultgren SJ, Larsson C, Almqvist F, Stallings CL. 2019. Chemical disarming of isoniazid resistance in *Mycobacterium tuberculosis*. Proc Natl Acad Sci U S A 116:10510–10517.

62. Lu P, Asseri AH, Kremer M, Maaskant J, Ummels R, Lill H, Bald D. 2018. The anti- mycobacterial activity of the cytochrome *bcc* inhibitor Q203 can be enhanced by small-molecule inhibition of cytochrome *bd*. Sci Rep 8:2625.

63. Beites T, O’Brien K, Tiwari D, Engelhart CA, Walters S, Andrews J, Yang HJ, Sutphen ML, Weiner DM, Dayao EK, Zimmerman M, Prideaux B, Desai PV, Masquelin T, Via LE, Dartois V, Boshoff HI, Barry CE, 3rd, Ehrt S, Schnappinger D. 2019. Plasticity of the Mycobacterium tuberculosis respiratory chain and its impact on tuberculosis drug development. Nat Commun 10:4970.

64. Lu X, Williams Z, Hards K, Yang J, Cheung CY, Aung HL, Wang B, Liu Z, Hu X, Lenaerts A, Woolhiser L, Hastings C, Zhang X, Wang Z, Rhee K, Ding K, Zhang T, Cook GM. 2019. Pyrazolo[1,5-a]pyridine inhibitor of the respiratory cytochrome *bcc* complex for the treatment of drug-resistant tuberculosis. ACS Infect Dis 5:239–249.

65. Pandey AK, Sassetti CM. 2008. Mycobacterial persistence requires the utilization of host cholesterol. Proc Natl Acad Sci U S A 105:4376–4380.

66. Pethe K, Sequeira PC, Agarwalla S, Rhee K, Kuhen K, Phong WY, Patel V, Beer D, Walker JR, Duraiswamy J, Jiricek J, Keller TH, Chatterjee A, Tan MP, Ujjini M, Rao SP, Camacho L, Bifani P, Mak PA, Ma I, Barnes SW, Chen Z, Plouffe D, Thayalan P, Ng SH, Au M, Lee BH, Tan BH, Ravindran S, Nanjundappa M, Lin X, Goh A, Lakshminarayana SB, Shoen C, Cynamon M, Kreiswirth B, Dartois V, Peters EC, Glynne R, Brenner S, Dick T. 2010. A chemical genetic screen in *Mycobacterium tuberculosis* identifies carbon-source-dependent growth inhibitors devoid of in vivo efficacy. Nat Commun 1:57.

67. Eoh H, Rhee KY. 2013. Multifunctional essentiality of succinate metabolism in adaptation to hypoxia in *Mycobacterium tuberculosis*. Proc Natl Acad Sci U S A 110:6554–9.

68. Hards K, Cook GM. 2018. Targeting bacterial energetics to produce new antimicrobials. Drug Resist Updat 36:1–12.

69. Kim JH, Wei JR, Wallach JB, Robbins RS, Rubin EJ, Schnappinger D. 2011. Protein inactivation in mycobacteria by controlled proteolysis and its application to deplete the beta subunit of RNA polymerase. Nucleic Acids Res 39:2210–20.

70. Kim JH, O’Brien KM, Sharma R, Boshoff HI, Rehren G, Chakraborty S, Wallach JB, Monteleone M, Wilson DJ, Aldrich CC, Barry CE, 3rd, Rhee KY, Ehrt S, Schnappinger D. 2013. A genetic strategy to identify targets for the development of drugs that prevent bacterial persistence. Proc Natl Acad Sci U S A 110:19095–100.

71. Lamprecht DA, Finin PM, Rahman MA, Cumming BM, Russell SL, Jonnala SR, Adamson JH, Steyn AJ. 2016. Turning the respiratory flexibility of *Mycobacterium tuberculosis* against itself. Nat Commun 7:12393.

72. Berube BJ, Russell D, Castro L, Choi SR, Narayanasamy P, Parish T. 2019. Novel MenA Inhibitors Are Bactericidal against *Mycobacterium tuberculosis* and Synergize with Electron Transport Chain Inhibitors. Antimicrob Agents Chemother 63.

73. Lu P, Heineke MH, Koul A, Andries K, Cook GM, Lill H, van Spanning R, Bald D. 2015. The cytochrome *bd*-type quinol oxidase is important for survival of *Mycobacterium smegmatis* under peroxide and antibiotic-induced stress. Sci Rep 5:10333.

74. Bald D, Villellas C, Lu P, Koul A. 2017. Targeting Energy Metabolism in *Mycobacterium tuberculosis*, a New Paradigm in Antimycobacterial Drug Discovery. mBio 8.

75. Sambandamurthy VK, Wang X, Chen B, Russell RG, Derrick S, Collins FM, Morris SL, Jacobs WR, Jr. 2002. A pantothenate auxotroph of *Mycobacterium tuberculosis* is highly attenuated and protects mice against tuberculosis. Nat Med 8:1171–4.

76. Berney M, Berney-Meyer L. 2017. Mycobacterium tuberculosis in the Face of Host-Imposed Nutrient Limitation. Microbiol Spectr 5.

77. Berney M, Berney-Meyer L, Wong KW, Chen B, Chen M, Kim J, Wang J, Harris D, Parkhill J, Chan J, Wang F, Jacobs WR, Jr. 2015. Essential roles of methionine and S-adenosylmethionine in the autarkic lifestyle of Mycobacterium tuberculosis. Proc Natl Acad Sci U S A 112:10008–13.

78. Lambert R, Pearson J. 2000. Susceptibility testing: accurate and reproducible minimum inhibitory concentration (MIC) and non-inhibitory concentration (NIC) values. J Appl Microbiol 88:784–790.

